# Signatures of copy number alterations in human cancer

**DOI:** 10.1101/2021.04.30.441940

**Authors:** Christopher D. Steele, Ammal Abbasi, S. M. Ashiqul Islam, Azhar Khandekar, Kerstin Haase, Shadi Hames, Maxime Tarabichi, Tom Lesluyes, Adrienne M. Flanagan, Fredrik Mertens, Peter Van Loo, Ludmil B. Alexandrov, Nischalan Pillay

**Affiliations:** Research Department of Pathology, Cancer Institute, University College London, London, WC1E 6BT, UK; Department of Cellular and Molecular Medicine, UC San Diego, La Jolla, CA, 92093, USA; Department of Bioengineering, UC San Diego, La Jolla, CA, 92093, USA; Moores Cancer Center, UC San Diego, La Jolla, CA, 92037, USA; Cancer Genomics Laboratory, The Francis Crick Institute, London, NW1 1AT, UK; Department of Cellular and Molecular Pathology, Royal National Orthopaedic Hospital NHS Trust, Stanmore, HA7 4LP, UK; Division of Clinical Genetics, Department of Laboratory Medicine, Lund University, Lund, Sweden; Department of Clinical Genetics and Pathology, Division of Laboratory Medicine, Lund, Sweden

**Author notes:** Denotes equal contributions. Correspondence and request for materials should be addressed to and.

## Abstract

The gains and losses of DNA that emerge as a consequence of mitotic errors and chromosomal instability are prevalent in cancer. These copy number alterations contribute to cancer initiaition, progression and therapeutic resistance. Here, we present a conceptual framework for examining the patterns of copy number alterations in human cancer using whole-genome sequencing, whole-exome sequencing, and SNP6 microarray data making it widely applicable to diverse datasets. Deploying this framework to 9,873 cancers representing 33 human cancer types from the TCGA project revealed a set of 19 copy number signatures that explain the copy number patterns of 93% of TCGA samples. 15 copy number signatures were attributed to biological processes of whole-genome doubling, aneuploidy, loss of heterozygosity, homologous recombination deficiency, and chromothripsis. The aetiology of four copy number signatures are unexplained and some cancer types have unique patterns of amplicon signatures associated with extrachromosomal DNA, disease-specific survival, and gains of proto-oncogenes such as *MDM2*. In contrast to base-scale mutational signatures, no copy number signature associated with known cancer risk factors. The results provide a foundation for exploring patterns of copy number changes in cancer genomes and synthesise the global landscape of copy number alterations in human cancer by revealing a diversity of mutational processes giving rise to copy number changes.

## MAIN

Genome instability is a hallmark of cancer leading to changes of the genomic DNA sequence, aneuploidy, and focal copy number alterations^1^. Both aneuploidy and sub-chromosomal copy number alterations have been previously associated with increased cell proliferation, poor prognosis, and reduced infiltration of immune cells^2–6^. Aneuploidy and genome-wide structural variation may originate from mitotic slippage, spindle multipolarity, and breakage-fusion-bridge (BFB) cycles^7^. Besides chromosome mis-segregation, other macroevolutionary mechanisms lead to changes in genomic copy number, including whole-genome doubling (WGD), where the entire chromosomal content of a cell is duplicated^8^ and chromothripsis where a “genomic catastrophe” leads to clustered rearrangements and oscillating copy number^9^. These evolutionary events may occur multiple times at different intensities during tumour development leading to a highly complex genome^10–12^.

The complex structural profiles of human cancers are mirrored by the intricate patterns of somatic mutations imprinted on cancer genomes at a single nucleotide level. Previously, we developed a computational framework that allows separating these intricate patterns of somatic mutations into individual mutational signatures of single base substitutions (SBS), doublet base substitutions (DBS), and small insertion or deletions (ID)^13,14^. Analyses of mutational signatures have provided unprecedented insights into the exogenous and endogenous processes moulding cancer genomes at a single nucleotide level with mutational signatures attributed to exposures to environmental mutagens, failure of DNA repair, infidelity/deficiency of polymerases, iatrogenic events, and many others^15–22^.

We recently developed a “mechanism-agnostic” approach for summarising allele-specific copy number patterns in whole genome sequenced sarcomas^23^ which we term copy number signatures. Other cancer subtype-specific methods for interrogating copy number patterns have been created and applied to ovarian cancer and breast cancer^24,25^. While these initial approaches have led to biological and clinical insights, there is currently no approach that allows interrogating copy number signatures across multiple cancer types and across different experimental assays.

To address this gap we developed a new framework for deciphering copy number signatures across cancer types and demonstrate its applicability to whole-genome sequencing, whole-exome sequencing, and SNP6 microarray data. We identified 19 distinct copy number signatures many of which are shared across multiple histologies and others that are specific to certain cancer subtypes. Extensive computational simulations, refinement and statistical association analyses were used both to assign processes to many of these signatures and to demonstrate their biological and clinical relevance. Overall, our findings shed light on the processes of chromosomal segregation errors and provide a method to distil the ensuant complex genomic configurations.

### A framework for pan-cancer classification of copy number alterations

We examined the allele-specific copy number profiles of 9,873 primary cancer samples across 33 cancer types from The Cancer Genome Atlas project (TCGA; **Supplementary Table 1**). The severity of genomic instability, measured by number of copy number segments, proportion of the genome displaying loss of heterozygosity (LOH) and genome doubling status vary greatly amongst cancer types (**Fig. 1*a-b***). Nevertheless, a linear relationship was observed between the number of segments and proportion of genomic LOH, varying from cancers with diploid and copy number “quiet” genomes (*e.g.,* acute myeloid leukaemia, thymoma, and thyroid carcinoma; **Fig. 1*a***) to cancers with highly aberrant copy number profiles (*e.g.,* ovarian carcinomas and sarcomas; **Supplementary Fig. 1*a-b*).** This linear relationship fails to hold only for adrenocortical carcinoma and chromophobe renal cell carcinoma both of which demonstrate enrichment of LOH without enrichment of copy number segmentation (**Supplementary Fig. 1*a-c***). Additionally, considerable variability of ploidy was observed both between and within cancer types (**Fig. 1*b***, **Supplementary Fig. 1*d***).

**Figure 1.**
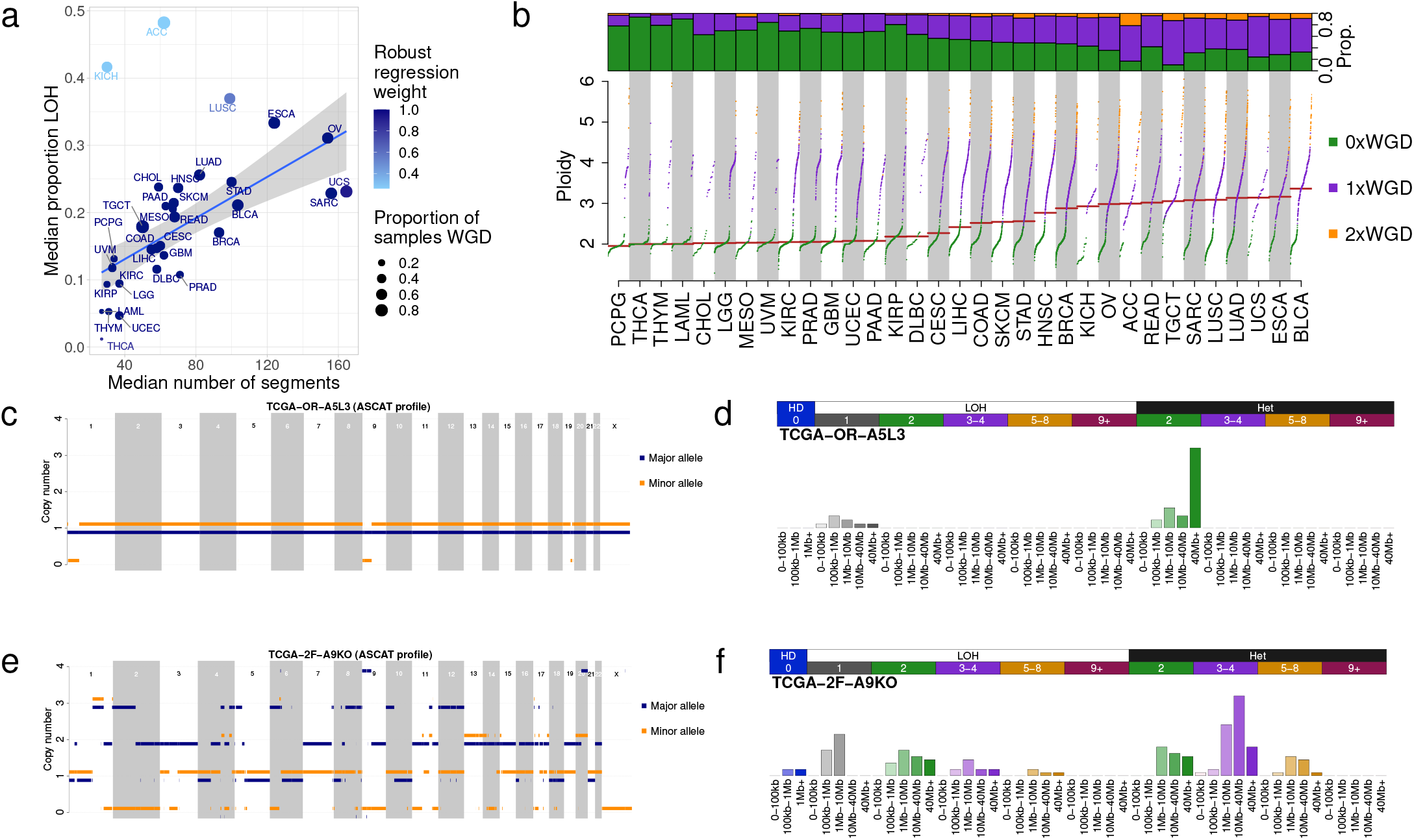
Pan-cancer copy number characteristics in TCGA. **a)** Copy number characteristics of 33 tumour types included in TCGA. Median number of segments in a copy number profile (x-axis), median proportion of the genome that has loss of heterozygosity (y-axis) and the proportion of samples that have undergone one or more whole genome doubling events (size). The line of best fit from a robust linear regression is shown, where the colour of points indicates the weight of the tumour type in the regression model. **b)** Ploidy characteristics of all TCGA samples, split by tumour type. Bottom panel: ploidy (y-axis) against quantile of ploidy (y-axis) for each sample in a tumour type, where samples are coloured by their genome doubling status: 0×WGD=non genome doubled (green), 1×WGD=genome doubled (purple), 2×WGD=twice genome doubled (orange). Top panel: proportion of samples in each tumour type that are 0, 1 or 2×WGD. **c)** Allele-specific copy number profile from a majority diploid sample (sample ID: TCGA-OR-A5L3, tumour type: ACC). Copy number (y-axis) across the genome (x-axis) is given for both the major (blue) and minor (orange) allele. **d)** Copy number summary for TCGA-OR-A5L3 after categorizing each of the segments. Segments are characterized first as homozygously deleted (left, blue), LOH (middle, white) or heterozygous (right, black), then by copy number states: TCN=0 (blue), TCN=1 (grey), TCN=3-4 (purple), TCN=5-8 (orange) and TCN=9+ (red). Finally, segments are categorized by segment size (increasing colour saturation indicates increasing segment size): 0-100kb, 100kb-1Mb, 1Mb-10Mb, 10Mb-10Mb and 40Mb+ (bottom labels). Homozygous deletions have a largest segment size category of 1Mb+. **e)** Allele-specific copy number profile for a highly aberrant sample (sample ID: TCGA-2F-A9KO, tumour type: BLCA). **f)** Copy number summary for TCGA-2F-A9KO.

To capture biologically relevant copy number features, we developed a classification framework that encodes the copy number profile of a sample by summarizing the counts of segments into a 48-dimensional vector. Specifically, copy number segments were classified into three heterozygosity states: heterozygous segments with copy number of {*A*>0, *B*>0} (numbers reflect the counts for major allele *A* and minor allele *B*), segments with LOH with copy number of {*A*>0, *B*=0}, and segments with homozygous deletions {*A*=0, *B*=0}. Segments were further subclassified into 5 classes based on the sum of major and minor allele (total copy number, TCN; **Supplementary Fig. 1*e***) and chosen for biological relevance: TCN=0 (homozygous deletion), TCN=1 (deletion leading to LOH), TCN=2 (wild type, including copy-neutral LOH), TCN=3 or 4 (minor gain), TCN=5 to 8 (moderate gain), and TCN>=9 (high-level amplification). Each of the heterozygous and LOH total copy numbers were then subclassified into five classes based on the size of their segments: 0 – 100kb, 100kb – 1Mb, 1Mb – 10Mb, 10Mb – 40Mb, and >40Mb (the largest category for homozygous deletions was restricted to >1Mb) in order to capture focal, large scale, and chromosomal copy number changes. The segment sizes were selected to ensure that a sufficient proportion of segments were classified in each category resulting in a reasonable representation across the pan-cancer TCGA dataset (**Supplementary Fig. 1*f-h***). Two examples, one encoding a mostly diploid adrenocortical carcinoma (**Fig. 1*c-d***) and another encoding a genomically unstable bladder cancer (**Fig. 1*e-f***), are provided to illustrate the classification framework.

To determine the generalizability of our framework for pan-cancer classification of copy number alterations across experimental platforms, we performed a comparative analysis of samples simultaneously profiled with SNP6 microarrays, whole-exome sequencing (282 samples), and whole-genome sequencing (512 samples).

Optimisation of the copy number calling strategy (**Methods**) resulted in remarkably similar profiles between distinct experimental assays. Specifically, copy number profiles derived from exome sequencing data had a median cosine similarity of 0.925 with copy number profiles derived from SNP6 microarrays (**Supplementary Fig. 1*i***). Copy number profiles derived from whole-genome sequencing data exhibited median cosine similarities of 0.933 and 0.852 with profiles derived from SNP6 microarrays or exome sequencing, respectively (**Supplementary Fig. 1*j-k***). These similarities are considerably better than similar comparisons observed for mutational signatures of single base substitutions derived from whole-genome and exome sequencing (median cosine similarity=0.55).

### The repertoire of copy number signatures in human cancer

Copy number profiles from SNP6 microarrays (n=9,873) were concatenated into cancer type-specific matrices and separately in a global pan-cancer matrix. These matrices were decomposed using our previously established approach^26^ for deriving a reference set of signatures (**Methods**). The approach allowed the identification of both the shared patterns of copy number across all examined samples, termed, *copy number signatures*, as well as the quantification of the number of segments attributed to each copy number signature in each sample, termed, *signature attribution*.

By applying our copy number signature framework (**Methods**) we identified 19 distinct pan-cancer signatures (**Fig. 2*a***; **Supplementary Table 2**). These signatures accurately explained the copy number profiles (p-value<0.05, Methods) of 93% of the examined TCGA samples. The remaining 7% were poorly explained due to a combination of a low number of segments and/or a high diversity of copy number states in the copy number profile or few operative signatures identified (**Supplementary Figs. 2*a-c***). The 19 signatures were categorized into 6 groups based on their most prevalent features. CN1 and CN2 are primarily defined by >40Mb heterozygous segments with total copy number (TCN) of 2 and 3-4 respectively. CN3 is characterized by heterozygous segments with sizes above 1Mb and TCN between 5 and 8. CN4-8 each have segment sizes between 100kb and 10Mb but with different TCN or LOH states. CN9-12 each have numerous LOH components with segment size <40Mb. CN13-14 have whole-arm or whole-chromosome scale LOH events (>40Mb). CN15 consists of LOH segments with TCN between 2 and 4 as well as heterozygous segments with TCN between 3 and 8, each with segment sizes 1-40Mb. CN16-19 exhibited complex patterns of copy number alterations that are uncommon but are seen in distinct cancer types. Additionally, 3 artefactual signatures (CN20-22) indicative of copy number profile over-segmentation were identified (**Supplementary Fig. 2*d***). To determine if the copy number signatures would generalize between platforms, we compared copy number signatures derived from whole-genome and whole-exome sequencing with SNP6 array signatures which showed a strong concordance with a median cosine similarity between signatures above 0.80 (**Supplementary Fig. 2*e-h***).

**Figure 2.**
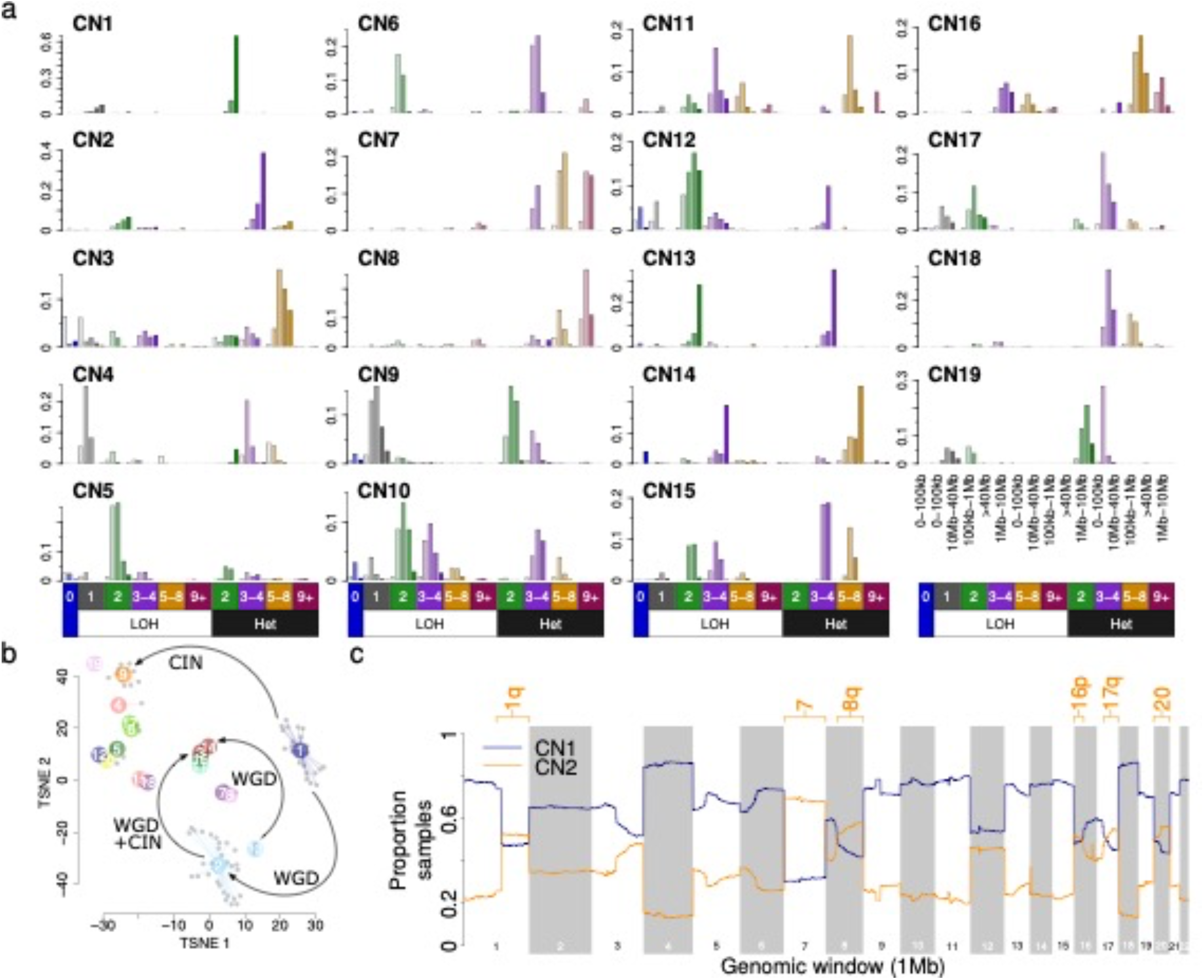
Patterns of pan-cancer copy number signatures. **a)** 19 identified non-artefactual copy number signatures in TCGA that are not linear combinations of any other. LOH status and total copy number are indicated below each column. Segment sizes for select bars are shown in the bottom right. Increasing saturation of colour indicates increasing segment size. **b)** TSNE representation of all non-artefactual consensus signatures (colours) and the individual signatures that were combined to form each consensus signature (grey). Inferences about the relationships between signatures (see Supplementary Figure 3) are indicated with arrows; WGD=whole-genome doubling, CIN=chromosomal instability. **c)** CN1 (blue) and CN2 (orange) recurrence (y-axis) across the genome (x-axis) in 472 highly aneuploid samples where CN1+CN2 attribution = 1. Chromosome arms with >50% samples attributed to CN2 are labelled.

### The transitional behaviour of copy number signatures

The catalogue of somatic mutations of a cancer genome is the cumulative result of the mutational processes that have been operative over the lifetime of the cell from which the cancer has derived^27^. Analysis of SBS and ID mutational signatures have used assumptions and prior evidence that individual mutations are independent and additive^28^. However, this assumption is clearly violated for large-scale macro-evolutionary events such as whole-genome doubling^29^.

We therefore generated several synergistic lines of evidence to investigate the impact of genome doubling on copy number signatures. First, each copy number signature was tested for enrichment in non-, once- or twice-genome doubled samples (**Supplementary Fig. 3*a-b***). Second, *in silico* simulations of genome doubling on the extracted signatures were performed (**Methods**; **Supplementary Fig. 3*c***). Third, copy number profiles arising from dynamics of whole-genome doubling and chromosomal instability (CIN) were simulated (**Supplementary Fig. 3*d***) and re-examined for the previously derived signatures (**Supplementary Fig. 3*e***).

By combining the preceding set of experiments, we revealed a transitional behaviour of copy number signatures with one signature being completely replaced by another upon genome doubling (**Fig. 2*b***). In this model, a cancer with a diploid signature (CN1), may undergo genome doubling, thus altering signature CN1 into signature CN2, or may undergo chromosomal instability transforming signature CN1 into signature CN9. Through a combination of CIN and genome doubling CN2 may also be changed to CN3. Additionally, CN13 and CN14 may be linked through genome doubling, on the background of early chromosomal losses.

While macro-evolutionary events have a transitional effect on copy number signatures, we hypothesized that smaller-scale events, such as segmental aneuploidy, may reflect an additive behaviour. To investigate this, we focused on the ploidy-associated signatures CN1 and CN2, where a combination of both signatures indicates a hyper-diploid or sub-tetraploid profile. Interestingly, each signature was found at below 50% attribution in approximately a quarter of TCGA samples, suggestive of potential aneuploidy in a considerable proportion of samples. We mapped these signatures across the cancer genomes with mixtures of attributions from signatures CN1 and CN2 (**Supplementary Fig. 3*f***). This analysis recapitulated known patterns of aneuploidy in human cancer^30,31^, including gains of chromosomes 1q, 7, 8q, 16p, 17q, and 20 in more than 50% of TCGA samples (**Fig. 2*c***).

### The landscape of copy number signatures

Next, we surveyed the distribution of the 19 signatures across the different cancer types (**Fig. 3**). Unsurprisingly, the ploidy associated signatures CN1 and CN2 were found in most samples across all cancer types with different median attributions.

**Figure 3.**
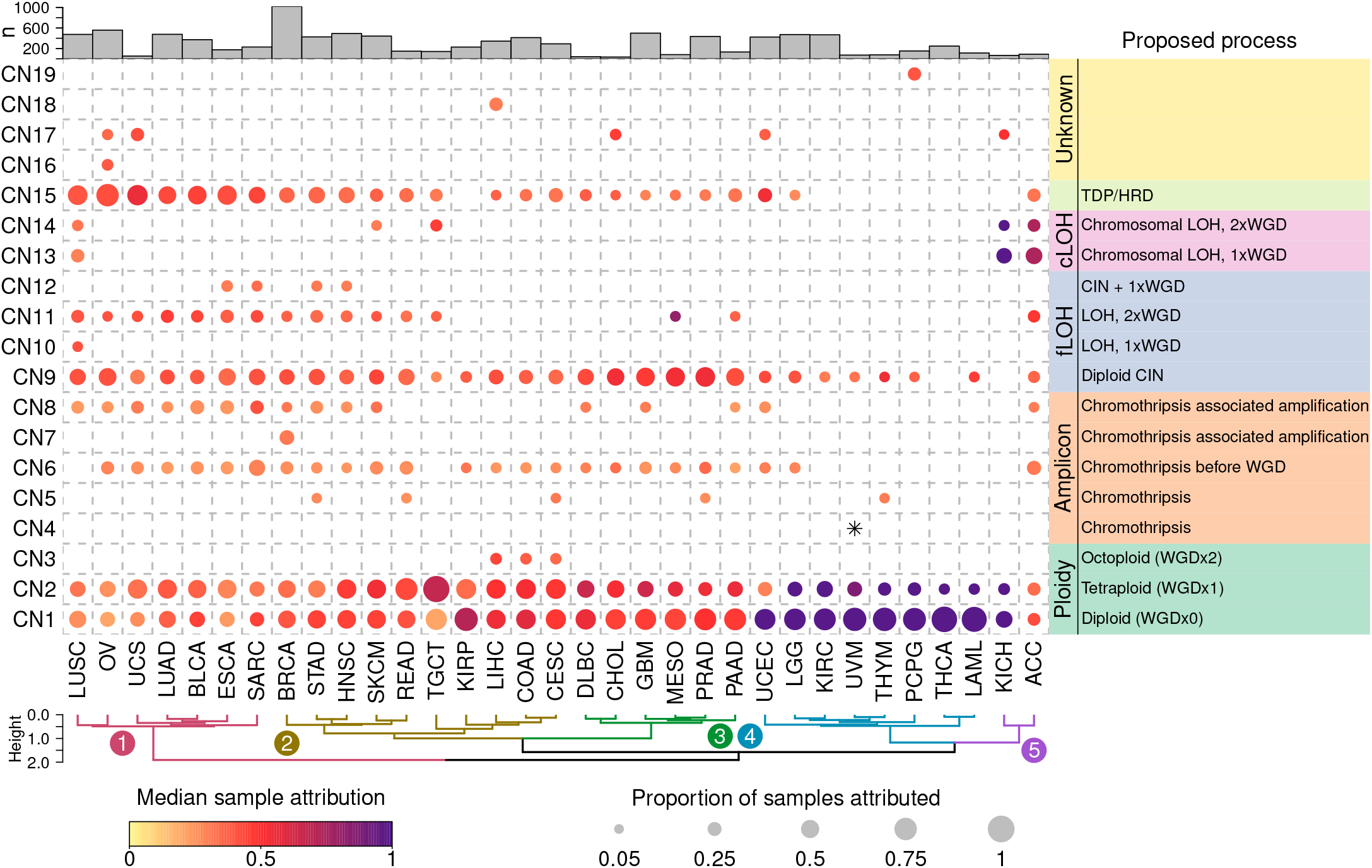
Distribution of copy-number signatures across human cancers. Attributions of the 19 non-artefactual signatures (y-axis) split by tumour type (x-axis), showing both the proportion of each tumour type exposed to each signature (size), and the median exposure of those tumours that are exposed to the signatures in each tumour type (colour). Tumour/signature combinations with less than 5% of samples exposed to the signature are not shown (except for CN4 in UVM, denoted with a *). Hierarchical clustering is shown below, sample sizes are shown above. Proposed processes are shown to the right.

Signatures CN4, CN7, CN10, CN16, CN18, and CN19 were derived through cancer type extractions and therefore unique to uveal melanoma, breast cancer, lung squamous carcinoma, ovarian carcinoma, liver cancer and paragangliomas, respectively. Signatures CN4-8 all showed segments of high total copy number and were seen in tumour types with known prevalent amplicon events^32^. CN9-CN12 showed differing patterns of hypodiploidy, LOH < 40Mb and WGD reflective of chromosomal instability. Signatures CN13 and CN14 were prevalent in adrenocortical carcinoma and chromophobe renal cell carcinoma, suggesting a link with the known patterns of chromosomal LOH (cLOH) seen in these cancers^33,34^. Signature CN15 was prevalent in tumour types previously described as being enriched in the tandem duplicator phenotype (TDP)^35^. Different cancer lineages clustered together based on the prevalence of signatures; namely TDP, whole-genome duplication, diploid chromosomal instability, simple diploidy, and chromosomal LOH (**Fig. 3**). This segregation of cancer types and their constituent signatures reflects the known distributions of genome doubling and aneuploidy in human cancer^3,36^.

### Copy number signatures associated with amplicons

Oncogene amplification has been associated with aggressive behaviour in cancer^32^, and can originate through the processes of BFB cycles and chromothripsis^12,37^. Reasoning that signatures with high levels of total copy number (CN4, CN5, CN6, CN7, and CN8) could associate with genomic amplification we correlated these signatures with known classes of amplicons^32,38^. All amplicon signatures were positively associated with one or more amplicon types (**Fig. 4*a***); CN8 was strongly associated with all four classes of amplicon, but most strongly with extra-chromosomal circular DNA amplicons (ecDNA).

**Figure 4.**
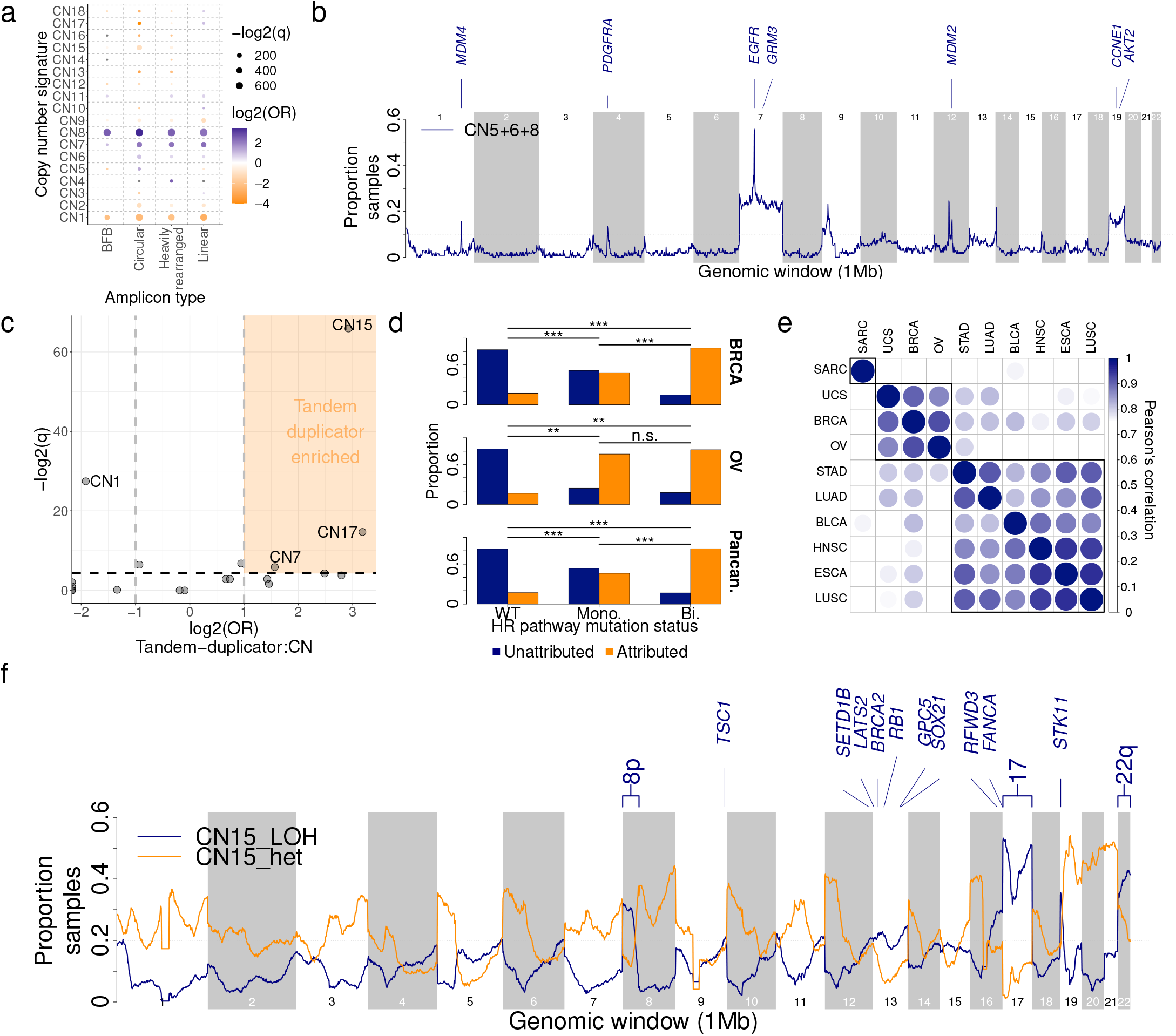
Biological inference of copy-number signatures. **a)** Associations between copy number signatures (y-axis) and amplicon structures (x-axis), displaying the q-value (size) and log2 odds ratio (colour) from a Fisher’s exact test of genomic regions attributed/not attributed to each signature against each amplicon type. Non-significant (q≥0.05) associations are not shown. BFB=breakage fusion bridge. CN8 was most strongly associated with circular amplicons: OR=10.8, q<5e-324. **b)** Recurrence of mapped amplicon signatures (CN5, CN6 and CN8) in 1Mb windows of the human genome across 134 GBM in which the amplicon signatures were attributed. Oncogenes in regions with >10% samples attributed to amplicon signatures are labelled. **c)** Associations between copy number signature attributed samples and tandem-duplicator phenotype samples, displaying −log2(q-values) (y-axis) and log2 odds ratios (x-axis). CN15 association: OR=7.6, q=1.5e-20, Fisher’s exact test. **d)** Correlation of CN15 attribution (y-axis) with mutational status of one or more genes of the homologous recombination pathway (x-axis) in breast cancer (top), ovarian cancer (middle) or pan-cancer (bottom). WT=wild type. Mono = Mono-allelic and Bi = bi-allelic. *=q<0.05, **=q<0.01, ***=q<0.001, n.s.=q≥0.05. **e)** Pearson’s correlation of recurrence of mapping of LOH segments of CN15 to the genome calculated for all pairwise comparisons of CN15-enriched tumour types. **f)** Recurrence of mapped CN15 in 1Mb windows of the human genome in all CN15 attributed BRCA, OV and UCS samples, split by LOH (blue) and heterozygous segments (orange). Tumour-suppressor genes in regions with >20% samples attributed to CN15 with LOH segments are labelled.

Recent evidence revealed that genomic amplification can evolve through interrelated processes of chromothripsis, BFB and ecDNA formation^11^. Therefore, we mapped the CN signatures with known regions of chromothripsis^39^ across the genome (**Methods**), revealing CN5-8 as being enriched in chromothriptic regions (**Supplementary Fig. 4*a***). Each of these signatures are dominated by small segments, while CN7-8 are both strongly associated with amplified chromothripsis^40^ (**Supplementary Fig. 4*b***) and complex chromothriptic events (**Supplementary Fig. 4*c***). Simulations of copy number profiles incorporating processes of chromothripsis, whole-genome doubling, and chromosomal duplication (**Supplementary Fig. 4*d***) demonstrated that CN4 to CN8 can be generated through chromothripsis-like events, and that these signatures reflect distinct life histories of tumours, such as chromothripsis before or after genome doubling (**Supplementary Figs. 4*c & e***).

Chromothripsis and gene amplification are both independently associated with poor prognosis^32,41^. Attribution of any of the five amplicon signatures in their respective cancer types resulted in a poor disease-specific survival in a univariate pan-cancer analysis (**Supplementary Fig. 5*a***). Similarly, multiple amplicon signatures were associated with a reduced disease-specific survival in multivariate pan-cancer and cancer type analyses with consistent results from analyses based on Cox-model hazard ratios (**Supplementary Fig. 5*b-c***) and analyses based on accelerated failure times (**Supplementary Fig. 5*d-e***). Cancer type-specific survival analysis revealed that patients with glioblastoma with operative signature CN5 had a poor disease-specific survival (172 days reduced median survival; **Supplementary Figure 5*d***). To determine the topographic localization of the amplification events, we mapped the amplicon signatures operative in glioblastoma (CN5, CN6, and CN8) across the genome which revealed recurrence of regions involving *EGFR, PDGFRA* and *MDM2* (**Fig. 4*b***) in keeping with previous reports of chromothripsis-associated amplification of these genes^42^.

### Copy number signatures associated with loss of heterozygosity

Loss of heterozygosity (LOH) is an important mechanism contributing to the inactivation of tumour suppressor genes during cancer development^39,43,44^. We found that 7 signatures positively correlated with LOH regions of the genome (**Supplementary Fig. 6*a***). Four of these signatures (CN9-12) were designated focal LOH (fLOH) signatures as they exhibited predominant segments sizes <40Mb (**Fig. 2**). The four fLOH signatures were recurrently found around tumour suppressor genes (**Supplementary Fig. 6*b***).

In adrenocortical carcinoma and chromophobe renal cell carcinoma a characteristic pattern of chromosome-level LOH leads to hypodiploidy^45,46^. We identified 2 signatures (CN13 and CN14) of chromosomal-scale LOH, each of which was enriched in both of these cancers (**Supplementary Fig. 6*c-d***). Mapping of these signatures to the genome revealed recurrent LOH in chromosome regions 1p, 3p, 5q, 9, 10q, 13q, and 17p (**Supplementary Fig. 6*e***), matching known patterns of aneuploidy in these tumours^33,34^ (**Supplementary Fig. 6f-g**).

### Copy number signature associated with tandem duplication and homologous recombination deficiency

Somatic tandem duplications (TD) are commonly found in breast and ovarian cancer^35,47,48^. Further, TD are strongly associated with failure of homologous recombination repair of DNA double strand breaks e.g. due to defective *BRCA1* or *BRCA2*^35,47,48^. A detailed characterization of TD across cancer has revealed three patterns with duplicated segments^35^ ranging around 10kb, 200kb, or 2Mb, respectively. CN15 has a segment size distribution that overlaps with the largest of these three patterns and was strongly associated with TD (**Fig. 4*c***, OR=7.6, q=1.5e-20, Fisher’s exact test) and enriched in cancer types known to show TD (**Supplementary Fig. 7*a***)^35^.

Consistent with prior observations for TD, an enrichment of CN15 is observed for samples harbouring mono-allelic defects in the homologous recombination pathway compared to wild-type samples for breast cancer (**Fig. 4*d***; OR=4.5 with q=6.1e-14; Fisher’s exact test), ovarian cancer (OR=15.3 with q=5.9e-3), and across all cancers (OR=4.2 with q=2.2e-106). Further enrichments of CN15 were observed in samples with bi-allelic defects in the homologous recombination pathway compared to samples with mono-allelic defects for breast cancer (**Fig. 4*d***; OR=6.2 with q=6.2e-5; Fisher’s exact test) and across all cancers (OR=5.7 with q=4.3e-16).

Prior analysis has shown that breakpoints resulting from TDs segregate non-randomly in the genome^35^. Mapping of CN15 to the genomes of CN15-enriched cancers revealed a tumour type-specific distribution of LOH segments (**Fig. 4*e***), but not of heterozygous segments (**Supplementary Fig. 7*b***). Breast and ovarian cancer as well as uterine carcinosarcoma displayed recurrent chromosomal LOH at 8p, 17 (including *BRCA1* and *TP53)*, and 22 (**Fig. 4*f***). Focal LOH was also observed on 9q around *TSC1*, 13q around *BRCA2* and *RB1*, and 19p around *STK11* (**Fig. 4*f***). In contrast CN15 attributed sarcomas display strong peaks of recurrent LOH around known sarcoma tumour suppressor genes^49^ (*CDKN2A*, *RB1*, and *TP53*; **Supplementary Fig. 7*c***). The 6 other tumour types enriched in CN15 display recurrent chromosomal LOH at 8p, 9p, 17p, 19p, and 21 (**Supplementary Fig. 7*d***).

### Copy number signatures associate with genomic features

To identify DNA damage repair mechanisms involved in the mutational processes giving rise to copy number signatures, we evaluated the associations between the activities of copy number signatures and single nucleotide level mutational signatures from both exome and whole genome sequencing data (**Fig. 5a).** As previously described SBS3 and ID6 are strongly associated with defective homologous recombination repair^14^. SBS2 and SBS13 are associated with APOBEC-mediated mutagenesis particularly seen near double stranded DNA breaks^50^. As expected, CN15 was strongly associated with SBS3 and ID6 derived from both WES and WGS data. Additionally, CN15 was associated with SBS2 and SBS13 providing a putative mechanistic link between APOBEC activity and CN15 in the context of TDPs. Negative associations were observed for diploid signature CN1 and APOBEC signatures SBS2 and SBS13 as well as for CN1 and tobacco-associated signature SBS4. These results indicate that diploid cancer genomes have lower APOBEC mutagenesis and that most cancers of tobacco smokers are not diploid.

**Figure 5.**
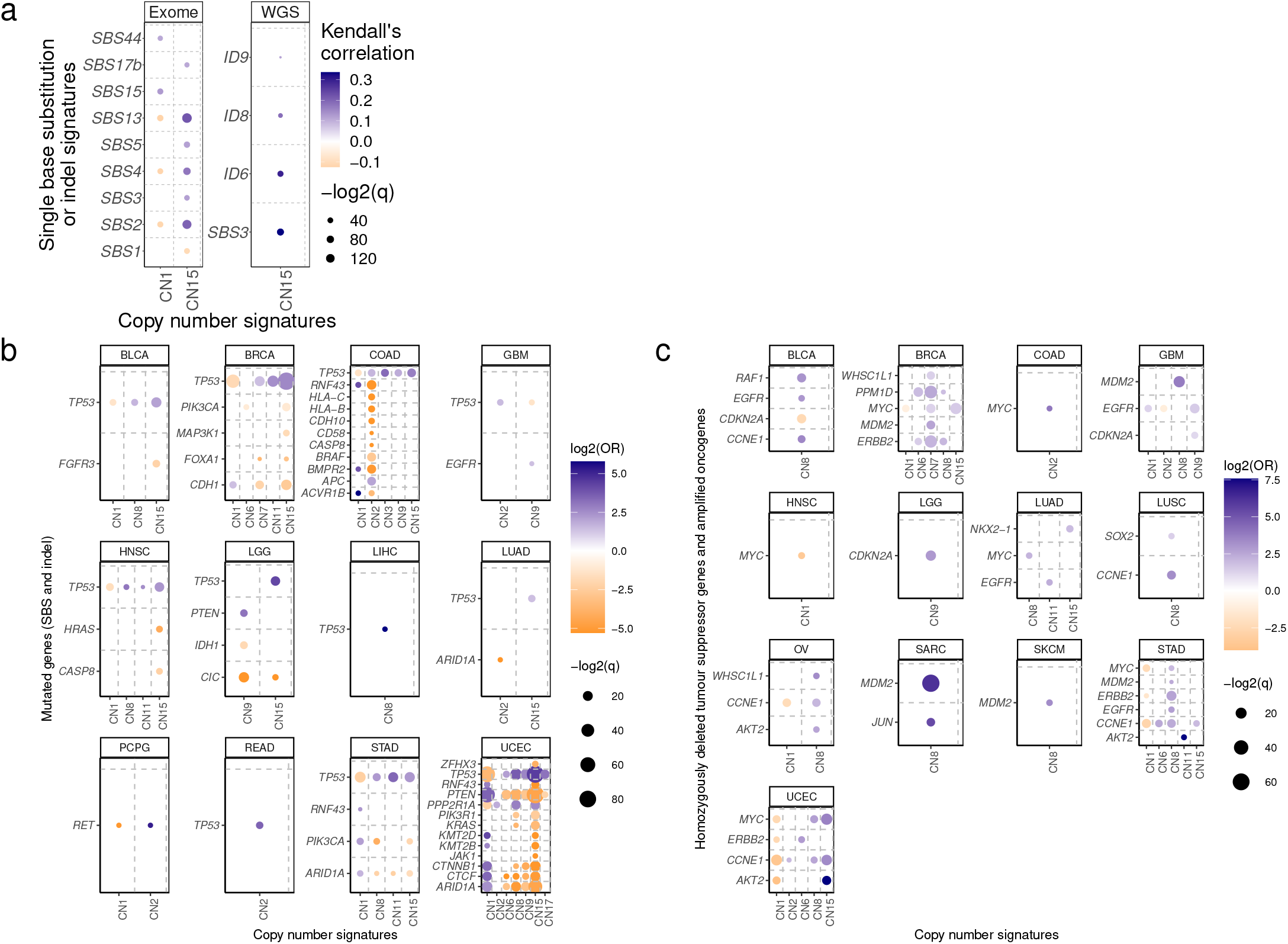
Genomic associations of copy number signatures. **a.** Correlation between copy number signature (y-axis) attribution and single base substitution signature (x-axis, SBS) exposure across TCGA exomes (left) and whole genomes (right). Strength of correlation is indicated by colour (orange=anti-correlated, blue=correlated), q-value is indicated by size of point. Only SBS signatures with any correlation between any copy number signatures with q<0.01 are shown. CN15 association with exome SBS3: Kendall’s correlation=0.12, q=7.5e-12. CN15 association with exome SBS2 and SBS13: Kendall’s correlation=0.2 and 0.22, q=1.6e-43 and 2.2e-50, respectively. CN15 association with WGS SBS3: Kendall’s correlation=0.34, q=1.1e-21. CN15 association with WGS ID6: Kendall’s correlation=0.29, q=4.7e-15. **b.** Associations between copy number signatures (x-axis) and driver gene SNV/indel status (y-axis) across each TCGA tumour type (panels). Effect size (log2 odds ratio, colour), and significance level (−log2 q-value, size) from a Fisher’s exact test are displayed. **c.** Associations between copy number signatures (x-axis) and driver gene copy number alteration status (y-axis, amplification for oncogenes, homozygous deletion for tumour-suppressor genes) across each TCGA tumour type (panels). Effect size (log2 odds ratio, colour), and significance level (−log2 q-value, size) from a Fisher’s exact test are displayed.

We next interrogated cancer driver gene mutations and copy number signatures and found significant differences between cancer types. A consistent finding across cancer was a positive association between *TP53* mutation and multiple copy number signatures (**Fig. 5*b***). *TP53* mutations were also associated with an increased diversity of copy number signatures (**Supplementary Fig. 8*a***; OR=3.42 with q=1.5e-49), supporting the link between *TP53* alteration and aneuploidy^3,51–53^. Mutations in *RNF43, HLA-B, HLA-C* and *BRAF* are commonly seen in microsatellite instable (MSI) colon cancers and were found to be negatively correlated with samples with tetraploid genomes (*i.e.,* CN2 attributed; **Supplementary Fig. 8*b***). MSI is associated with high immune cell infiltration whilst aneuploidy is associated with a decrease in leucocyte fraction^54^. Across multiple cancer types, we observe a general trend of decreased leucocyte fractions in cancers with copy number signatures of aneuploidy compared to diploid cancers (CN1; **Supplementary Fig. 8*c***). Similar to colon cancer, multiple cancer driver genes were associated with CN1/CN2 in endometrial cancer, largely driven by differential copy number and mutation patterns seen in microsatellite stable and unstable tumours (**Supplementary Fig. 8*d***).

To assess the relationships between copy number signatures and copy number driver genes, we evaluated the associations between attributions of copy number signatures and homozygous deletions of COSMIC tumour suppressor genes as well as between attributions of copy number signatures and amplifications of known proto-oncogenes^55^. Copy number drivers such as *MDM2, EGFR, CCNE1, MYC*, and *ERBB2* were strongly positively associated with amplicon signatures CN6-8 as well as CN15 (**Fig. 5*c***). In contrast, *CDKN2A* was the only homozygously deleted tumour suppressor gene associated with any signature, most commonly CN9.

In contrast to single-nucleotide level SBS and ID signatures^14^, no associations were found between any copy number signature and cancer risk factors: gender, smoking status, or alcohol consumption (**Supplementary Fig. 8*e***). Significant associations were found between age and copy number signature attribution in individual tumour types (**Supplementary Fig 8*f***), however, these were driven by tumour sub-type differences: serous *versus* endometrioid endometrial cancers (difference in mean age at diagnosis=4.7 years, p=9.0e-5, Mann-Whitney test) in which non-endometrioid endometrial cancers are strongly associated with HRD^56^ and enriched in CN15 (OR=16.7, p<7.1e-26, Fisher’s exact test); synovial sarcoma *versus* other sarcoma (difference in mean age at diagnosis=-22.3 years, p=4.3e-3, Mann-Whitney test) in which synovial sarcomas are karyotypically simple^49^ and enriched in CN1 (OR=Inf, p=2.3e-5, Fisher’s exact test).

## DISCUSSION

In this report, we provide the first pan-cancer framework for analysing copy number signatures as well as the first comprehensive analysis of copy number signatures in human cancer. The results revealed multiple distinct copy number signatures including ones attributed to ploidy, amplification, loss of heterozygosity, chromothripsis, and tandem duplications. Multiple signatures of unknown processes, cancer subtype specific signatures as well as artefactual signatures were identified. Unlike SBS and ID mutational signatures, copy number signatures did not associate with known cancer risk factors. Rather, copy number signatures reflect the activity of endogenous mutational processes such as homologous recombination deficiency, aberrant mitotic DNA replication, and chromothripsis^11,12^.

The field of copy number signatures is nascent, with three distinct methods previously implemented in three distinct tumour types^23–25^. As the field matures it will become increasingly clear which models are better suited to addressing specific clinical or biological questions. To resolve these questions, pan-cancer analyses utilizing all of these methods will be key, and we present here the first step towards that goal; a mechanism-agnostic pan-cancer compendium of allele-specific copy number signatures.

## ACKNOWLEDGEMENTS

NP is a Cancer Research UK Clinician Scientist fellow (Award - 18387). CDS undertook this work with support from Cancer Research UK Travel Award (Award no-27969). Support was provided to NP and AMF by the National Institute for Health Research, the University College London Hospitals Biomedical Research Centre, and the Cancer Research UK University College London Experimental Cancer Medicine Centre.

Alexandrov laboratory was supported by US National Institute of Health’s R01 ES030993 and R01 ES032547. LBA is an Abeloff V Scholar and he is supported by an Alfred P. Sloan Research Fellowship. Research at UC San Diego was also supported by a Packard Fellowship for Science and Engineering to LBA.

This work was supported by the Francis Crick Institute, which receives its core funding from Cancer Research UK (FC001202), the UK Medical Research Council (FC001202), and the Wellcome Trust (FC001202). For the purpose of Open Access, the authors have applied a CC BY public copyright licence to any Author Accepted Manuscript version arising from this submission. This project was enabled through access to the MRC eMedLab Medical Bioinformatics infrastructure, supported by the Medical Research Council (grant number MR/L016311/1). PVL is a Winton Group Leader in recognition of the Winton Charitable Foundation’s support towards the establishment of The Francis Crick Institute.

Compute resources were provided by UC San Diego through the Triton Shared Computing Cluster, and by UCL through the Myriad computing cluster.

The results shown here are in whole or part based upon data generated by the TCGA Research Network: https://www.cancer.gov/tcga.

Thanks to Dr Marnix Jansen and Dr Hamzeh Kayhanian for critical input to the work shown here.

## AUTHORS CONTRIBUTIONS

Study was conceived and designed by CDS, NP and LBA. Data analysis was performed by CDS, AA, SMAI, AK, KH, SH, MT and TL. Manuscript was written by CDA, NP and LBA. Interpretation of data and contributions to writeup were provided by MT, TL, AMF, FM and PVL.

## DATA AVAILABILITY

No new data was generated for this study. ASCAT copy number profiles that were generated for a different study and analysed here can be found at: https://github.com/VanLoo-lab/ascat/tree/master/ReleasedData/TCGA_SNP6_hg19

## CODE AVAILABILITY

Code for summarising copy number profiles into 48-length vectors can be found at: https://github.com/AlexandrovLab/SigProfilerMatrixGenerator

Code for extracting copy number signature can be found at: https://github.com/AlexandrovLab/SigProfilerExtractor

Code for decomposing copy number summaries into known copy number signatures can be found at: https://github.com/AlexandrovLab/SigProfilerSingleSample

Bespoke scripts for all other analysis are available from authors upon request.

## ONLINE METHODS

### Utilized datasets

Using SNP6 microarray data, copy number profiles were generated for 9,873 cancers and matching germline DNA of 33 different types from The Cancer Genome Atlas (TCGA)^43^ using allele-specific copy number analysis of tumours (ASCAT)^58^ with a segmentation penalty of 70 (**Supplementary Table 1**). Additionally, a set of whole-genome sequences from 512 cancers of the International Cancer Genome Consortium (ICGC) that overlapped with tumour profiles in TCGA were analysed^39^ to generate WGS-derived copy number profiles(see below). Lastly, a set of whole-exome sequences from 282 cancers from TCGA was analysed to generate exome-derived copy number profiles (see below).

### Copy number profile summarization

Copy number segments were categorized into three heterozygosity states: heterozygous (CN={>0,>0} for the major and minor alleles respectively), loss of heterozygosity (LOH; CN={>0,0}) and homozygous deletion (CN={0,0}). Segments were further subclassified into 5 categories of total copy number: CN0 reflects homozygous deletions, CN1 represents a genomic deletion, CN2 represents a diploid state, CN3-4 is a tri-to-tetraploid or gained state, CN5-8 is a penta-to-octoploid state and CN9+ represents high-level amplifications. Segments were further subclassified into 5 size categories: 0-100kb, 100kb-1Mb, 1Mb-10Mb, 10Mb-40Mb, and >40Mb. For homozygous deletions only 3 size categories were used: 0-100kb, 100kb-1Mb, and >1Mb. In this way copy number profiles were summarized as counts of 48 combined copy number categories defined by heterozygosity, copy number and size, which we will define as *N* = [*n*_1_, *n*_2_,…, *n*_48_]. For a given dataset, the copy number profiles of a set with *S* samples are then summarized as a nonnegative matrix with *S* × 48 dimensions.

### Deciphering signatures of copy number alterations

Copy number signatures were extracted by applying our previously developed approach for creating a reference set of signatures^14^. Specifically, SigProfilerExtractor v1.0.17^26^ was applied to the matrix encompassing all TCGA samples as well as separately to each matrix corresponding to an individual tumour type. In brief, SigProfilerExtractor utilizes nonnegative matrix factorization (NMF) to find a set of copy number signatures ranging from 1 to 25 components for each examined matrix. For each number of components, 250 NMF replicates with distinct initializations of the lower dimension matrices were performed on the Poisson resampled data. SigProfilerExtractor was used with default parameters, except for the initializations of the lower dimension matrices where random initialization was utilized consistent with our prior analyses of mutational signatures^14,59^ After performing 250 nonnegative matrix factorizations, SigProfilerExtractor clusters the factorization within each decomposition to automatically identify the optimum number of operative signatures that best explain the data without overfitting these data^26^.

As previously done^60^, the sets of all identified copy number signatures were combined into a reference set of pan-cancer copy number signatures by leveraging hierarchical clustering based on the cosine dissimilarities between each signature. The number of combined signatures is chosen to maximise the minimum average cosine similarity between each signature in a cluster and the mean of all samples in that cluster, to ensure that each copy number signature in a cluster has a high similarity to the combined copy number signature for that cluster. Simultaneously, the maximum cosine similarity between mean copy number signatures for each cluster is minimized, to ensure that each combined signature is distinct from all others. To avoid reference signatures being linear combinations of two or more other signatures, for each identified signature, a synthetic sample was created with the pattern of the signature multiplied by 1,000 copy number segments. Further, the synthetic sample was resampled with probabilities 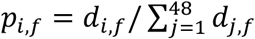 where *d_i,f_* is the strength of the *i*^th^ copy number category in the *f*^th^ identified signature. Each resampling was then scanned for activity of all other signatures from the reference set. If a resampled sample can be reconstituted with a cosine similarity >0.95 by 3 or fewer other signatures, the signature used to create the synthetic sample was deemed to be a linear combination of those signatures, and the signature was removed from the global reference set of signatures.

### Reference set of copy number signatures

Initially 28 pan-cancer copy number signatures were derived from the different SigProfilerExtractor analyses of the 9,873 copy number profiles from SNP microarrays. *In silico* evaluation and manual curation showed that 10 copy number signatures were linear combinations of two or more other signatures. Additionally, 3 signatures were deemed to be artefactual due to over-segmentation of copy number profiles. These artefactual signatures were removed from further analyses, as were the samples with any attribution of any of these artefactual signatures (116 samples; 1.2% of all TCGA samples). Moreover, samples with >25Mb of homozygous deletions across the genome were removed from downstream analysis (58 samples), leaving 9,699 samples for full analysis. Upon signature assignment (see below) 3 of the signatures that were removed due to linear combination were re-extracted within tumour-type specific assignment (cosine similarity=1), suggesting some copy number profiles could not be explained well without these 3 signatures. As a result, these 3 signatures were reintroduced into the compendium of signatures, leaving a total of 19 non-artefactual pan-cancer signatures of copy number alteration.

CN1-3 form a group of ploidy-associated signatures. CN1 and CN2 display TCN between 2 and 3-4 respectively, with predominantly >40Mb heterozygous segments. CN3 consists of predominantly heterozygous segments of TCN 5-8 with sizes >1Mb.

CN4-8 form a group of amplicon-associated signatures, that all have segment sizes predominantly between 100kb and 10Mb but with differing TCN or LOH states. CN4 consists of a mixture of LOH segments with TCN 1 and heterozygous segments with TCN 3-4. CN5 consists almost entirely of LOH segments with TCN 2. CN6 consists of a mixture of LOH segments with TCN 2 and heterozygous segments with TCN 3-4. CN7 consists of a mixture of heterozygous segments with TCN of 3-4, 5-8 and 9+.

CN8 consists of predominantly heterozygous segments with TCN 9+.

CN9-12 form a group of signatures with considerable LOH components. CN9 consists of a mixture of LOH segments with TCN 2 and heterozygous segments with TCN 2, each ranging from 100kb-40Mb. CN10 consists of a mixture of LOH segments with TCN 2 and 3-4 as well as heterozygous segments with TCN 3-4 between 100kb and 40Mb. CN11 consists of a mixture of LOH segments with TCN 3-4 and heterozygous segments with TCN 5-8, each at predominantly 1-10Mb. CN12 consists of mostly LOH segments of TCN 2 with sizes above 100kb and additional heterozygous segments of TCN 3-4 with sizes between 10 and 40Mb.

CN13-14 form a group of signatures with whole-arm or whole-chromosome scale LOH events. CN13 consists of LOH segments with TCN 2 and heterozygous segments with TCN 3-4, each at >40Mb, while CN14 is similar but with TCN 3-4 and 5-8 for LOH and heterozygous segments respectively.

CN15 has been associated with the tandem duplicator phenotype (**Fig. 4**). This signature consists of LOH segments of TCN 2 and 3-4 as well as heterozygous segments of TCN 3-4 and 5-8, each with segment sizes 1-40Mb.

CN16-19 originate from unknwon processes and are diverse in their copy number patterns. CN16 consists of predominantly heterozygous segments of TCN 4-8 at >1Mb, but with appreciable contributions of LOH segments with TCN 3-4 at >1Mb and heterozygous segments with TCN 9+ at >100kb. CN17 consists of segments between 100kb and 40Mb that are heterozygous with TCN 3-4 or less commonly LOH with TCN 1 or 2. CN18 consists of predominantly heterozygous segments with TCN 3-4 at 100kb-40Mb with some heterozygous segments of TCN 3-4 at 100kb-10Mb. CN19 consists of heterozygous segments with TCN 2 at >1Mb and many heterozygous segments with TCN 3-4 at 100kb-1Mb.

### Assignment of copy number signatures to individual cancer samples

The global reference set of copy number signatures was used to assign an activity for each signature to each of 9,873 examined samples using the decomposition module of the SigProfilerExtractor^26^. For the assignment, the information of the *de novo* signature and their activities assigned to each sample were used to implement the decomposition module with default parameters except for the NNLS addition penalty (*nnls_add_penalty*) which was set to 0.1, the NNLS removal penalty (*nnls_remove_penalty*) which was set to 0.01, and the initial removal penalty (*initial_remove_penalty*) which was set to 0.05. Signatures were assigned to samples in both tumour-specific evaluations and in a pan-cancer evaluation. As previously done^60^, the signature attributions from either tumour-specific or pan-cancer evaluations that gave the best cosine similarity between the input sample vector and the reconstructed sample vector were used as the attributions for that sample in all subsequent analyses.

### Copy number signatured derived from whole-genome and exome sequencing data

A set of samples from TCGA with both SNP-array and exome sequencing data were selected (*n*=282). Copy number profiles were generated from the exome sequencing data using ASCAT across all of the dbSNP common SNP positions with a segmentation penalty ranging from 20 to 140. Signatures were re-extracted for these 282 samples from both the SNP-array derived copy number profiles and the exome-derived copy number profiles, and the resulting signatures were compared.

For whole-genome sequencing data, we examined 512 whole-genome sequenced samples from the PCAWG project overlapping with TCGA samples with microarray data. Copy number profiles from whole-genome sequencing data were generated using ASCAT across the SNP6 positions, with a segmentation penalty ranging from 20 to 120. Signatures were extracted for samples with both SNP6 microarray derived copy number profiles and the WGS derived copy number profiles, and the extracted signatures were compared. In all cases, segmentation penalty of 70 gave the best concordance for both copy number profiles and extracted copy number signatures based on SNP6 microarray, whole-genome sequencing, and whole-exome sequencing data.

### Mapping copy number signatures to the landscapes of cancer genomes

Given the original copy number profiles, the identified signature matrix of *c* copy number classes by *f* signatures, and the signature activity matrix of *s* samples by *f* signatures, it is then possible to map signatures to the genomic landscape for each cancer sample. The probability of each copy number class, ***c***, having originated from each signature, ***i*** from a total of ***I*** signatures, in a sample ***j*** can be defined as:

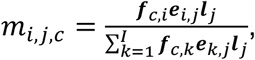

where ***f*** is the normalised signature matrix, ***e*** is the normalized attribution matrix, and ***l*** is a matrix of the number of segments in the copy number profile of each sample. The likelihood of each signature contributing to a given genomic window, here taken as each chromosome, is then the sum of copy number class probabilities for each segment in that window:

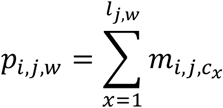

Once these chromosome likelihoods have been calculated, the individual segments in a chromosome are assigned to their maximum likelihood signature. Once copy number signatures have been mapped to the genome at a segment level, it is possible to interrogate the recurrence of signatures across the genome for a given set of copy number profiles. To do this, the genome is binned into 1Mb tiled windows. Within each window, the number of samples with a segment of a given copy number signature that overlaps the window is computed. This is repeated for each signature in each window.

### Associations between copy number signatures and events defined by genomic region

Localised events (chromothripsis^39^ and amplicon structure^38^) identified using WGS data were associated with mapped copy number signatures from TCGA for all available matching samples (chromothripsis *n*=657; amplicon *n*=1703). Each segment in every sample was categorised as overlapping or non-overlapping of a localized event. For each copy number signature, the association was then tested using a two-sided Fisher’s exact test on a contingency table of segments categorized as overlapping or non-overlapping of a localized event and assigned to or not assigned to the given copy number signature, across all samples. Multiple-testing correction was performed using the Benjamini-Hochberg method.

### Genome doubled copy number signatures

With the copy number categories being defined as 0, 1, 2, 3-4, 5-8, and 9+, it is possible to artificially ‘genome double’ any copy number category, other than 0, by assigning it to the next highest copy number category. In this way we artificially ‘genome doubled’ each signature by assigning the count for each copy number class to its next highest copy number class. First, the copy number 1 class is assigned a count of 0, then each copy number class is assigned the count of the preceding copy number class. For example, copy number class of 2 is assigned to the previous copy number class of 1, 3-4 assigned previous 2, *etc.*, until finally the copy number 9+ class is assigned a count that is the sum of the previous copy number 5-8 class and 9+ class. During this conversion, LOH and size categories are retained, so that the only shift is in copy number. Having performed this conversion, cosine similarities between the artificially ‘genome doubled’ signatures and the original signatures were calculated. Any genome-doubled and original signature pair that had a cosine similarity >0.85 was considered to contain a pair of signatures with analogous copy number patterns distinguished only by their genome doubling status.

### Associations between copy number signatures and ploidy

Ploidy for each copy number profile was calculated as the relative length weighted sum of total copy number across a sample. The proportions of the genome that displayed LOH (pLOH) were also calculated. Samples with a ploidy above −3/2*pLOH+3, meaning an LOH-adjusted ploidy of 3 or greater were deemed to be genome doubled samples, while samples with a ploidy above −5/2*pLOH+5, meaning an LOH-adjusted ploidy of 5 or greater, were deemed to be twice genome doubled samples. All other samples were considered as non-genome doubled samples. Each signature (CN1-19) was associated with each genome doubling category (GD×0, GD×1, and GD×2) using a one-sided Fisher’s exact test on a contingency table with samples categorized by whether the samples have >0.05 attribution to the given copy number signature or not, and whether the sample has the given genome doubled category or not. All p-values were corrected for multiple hypothesis testing using the Benjamini-Hochberg method.

### Associations between copy number signatures and known cancer risk factors

Associations between attributions of copy number signatures and attributions of single-base substitutions, indels, and doublet base signature exposures^14^ were performed using Kendall’s rank correlation. Only the significant associations found in both cancer-type specific and pan-cancer analysis were reported. For the cancer risk association analyses, copy number signatures were associated with gender^61^, tobacco smoking^18^, and alcohol drinking status^62^. For each copy number signature, the association was conducted using a two-sided Fisher’s exact test on a contingency table of a clinical feature categorized as present or absent and assigned to or not assigned to the given copy number signature across all samples. All p-values were corrected for multiple hypothesis testing using the Benjamini-Hochberg method.

Associations between copy number signature attribution (binarized to present or absent) and the tandem duplicator phenotype (also binarized to present or absent)^35^ were performed using a two-sided Fisher’s exact test (*n*=882). This was performed for each copy number signature separately. All p-values were corrected for multiple hypothesis testing using the Benjamini-Hochberg method and only associations with q<0.05 were reported.

Associations between copy number signature attribution (binarized to present or absent) and driver gene SNV/indel mutation status^63^ were performed within tumour types using a two-sided Fisher’s exact test (*n*=6,543 across all cancer types). This was performed for all copy number signature/gene combinations for which the gene was mutated in the given cancer type and the copy number signature was observed in the given cancer type. All p-values were corrected for multiple hypothesis testing using the Benjamini-Hochberg method and only associations with both q<0.05 and |log2(OR)|>1 were reported.

Driver copy number alterations of COSMIC cancer gene census genes^55^ were defined as: *(i)* homozygous deletion (CN={0,0}) of genes listed as deleted (D) in COSMIC mutation types; or *(ii)* amplification (CN>2*ploidy+1) of genes listed as amplified (A) in COSMIC mutation types. Associations were then performed on copy number driver alterations for SNV/indel driver gene alterations as above (*n*=9,699 across all cancer types).

The diversity of copy number signatures, as defined by Shannon’s diversity index, was associated with both SNV/indel and copy number driver gene mutations using a logistic regression model with binary diversity {>0, =0} as the dependent variable, and tumour type and gene mutation status as independent variables. LGG was taken as the reference tumour type. Only driver genes with >250 mutant samples in the dataset were included in the model.

Associations between copy number signature attribution (binarized to present or absent) and age at diagnosis (binarized to above or below median separately for each cancer type) were performed within cancer types using a two-sided Fisher’s exact test (*n*=8,841 across all cancer types). All p-values were corrected for multiple hypothesis testing using the Benjamini-Hochberg method and only associations with both q<0.05 and |log2(OR)|>1 were reported.

### Copy number signatures and defective homologous recombination

Signatures were tested for enrichment in tumour types using one-sided Mann-Whitney tests of signature attribution in a given tumour type versus all other tumour types. This was performed for all signature and tumour combinations. All p-values were corrected for multiple hypothesis testing using the Benjamini-Hochberg method.

Core homologous recombination (HR) repair pathway member genes were chosen to interrogate: *BRCA1*, *BRCA2*, *RAD51C*, *PALB2*^64,65^. Copy number alterations across these genes were identified based on ASCAT copy number profiles for homozygous deletions (*i.e.*, CN={0, 0}) and LOH (*i.e.*, CN={>0, 0}). Somatic SNVs and indels were taken from Ref. ^63^. Pathogenic germline variants in *BRCA1* and *BRCA2* were taken from Ref. ^66^. Samples were deemed as bi-allelically mutated for the HR pathway if homozygously deleted (HD) or if >1 of any of the other classes of alteration were present within any of the HR pathway genes. Mono-allelic loss was defined as 1 of any of the non-HD alterations within any of the HR pathway genes. Wildtype was defined as no alterations in any HR pathway genes. The associations between HR pathway status and CN15 were then restricted to only breast (*n*=589), ovarian (*n*=309), and pan-cancer (*n*=4,919). Two-sided fisher’s exact tests were performed between wild-type and mono-allelic samples, between wild-type and bi-allelic samples, and between mono-allelic and bi-allelic HR pathway status samples. All p-values were corrected for multiple hypothesis testing using the Benjamini-Hochberg method.

### Copy number signatures associated with changes of overall survival

Survival data for 11,160 TCGA patients were obtained from the TCGA Clinical data Resource R package^67^. Univariate disease specific survival analysis for signatures was performed using a log-rank test and Kaplan-Meier curves in R, with groups being unattributed (attribution=0) and attributed (attribution>0) for each signature separately, or for summed attributions of a set of signatures (*e.g.*, amplicon signatures).

Multivariate disease-specific survival analysis was performed using the Cox’s proportional hazards model in R with Boolean attributed/non-attributed variables for each copy number signature and tumour type as covariates. To account for potential violations of Cox’s model’s proportional hazards assumption, we also conducted the same analysis using the accelerated failure time model with the Weibull distribution using the flexsurvreg function in R. All p-values were corrected for multiple hypothesis testing using the Benjamini-Hochberg method.

### Simulating copy number profiles

#### Simulation framework

Genomes were initialized as 23 pairs of individual chromosomes, with lengths corresponding to those seen in the human genome, where the 23^rd^ pair could be either *X, X* or *X, Y*. Each chromosome was initialized as a data table with chromosome (1-22, X, Y), start position, end position, and allele (either A or B). Genomic events were recorded as altering one of these data tables in the appropriate way, adding or removing segments as necessary. Gains and losses: The log_10_(size) of sub-chromosomal gains were drawn from a Gaussian mixture with components:

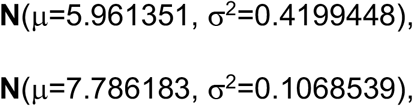

at proportions p_1_=0.7360366 and p_2_=1-p_1_. The log_10_(size) of sub-chromosomal losses were drawn from a gaussian mixture with components:

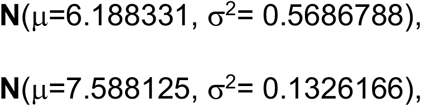

at proportions p_1_=0.6472512 and p_2_=1-p_1_. The parameters for the various distributions were estimated from samples in TCGA that were predominantly diploid (CN1+CN9 attribution>0.8) from segments that were copy number 1 for the loss distributions, and copy number 3 for the gain distributions. Parameters were estimated using a Gaussian mixture model on the log_10_(sizes) of the appropriate segments with two components due to the bimodal nature of the segment length distributions.

First the chromosome on which the gain/loss will occur is randomly sampled with probabilities 1/*n*, where *n* is the number of separate chromosomes in the current genome. The event size, λ, is then drawn from the previously stated multinormal distributions; if an event size greater than the chromosomal size is drawn, then a new size is drawn. The start of the event, *b_1_*, is then drawn from a uniform distribution,

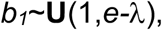

where *e* is the cumulative length of the chosen chromosome, and the end of the event, *b_2_*=*b_1_*+λ.

Gains are treated as tandem duplications, so that the gained region is inserted immediately after the start breakpoint. On unaltered chromosome, this will alter the chromosome from a single segment with start=1 and end=*e* to a chromosome with four segments, with starts=[1,*b_1_+1*,*b_1_+1*,*b_2_+1*] and ends=[*b_1_*,*b_2_*,*b_2_*,*e*], each with the chosen chromosome identity and allele; note that this will eventually lead to a copy number profile with 3 segments with starts==[1,*b_1_+1*,*b_2_+1*] and ends=[*b_1_*,*b_2_*,*e*]. A loss will instead lead to a chromosome with two segments with starts=[1,*b_2_*] and ends=[*b_1_*,*e*].

#### Simulating chromothripsis

For chromothriptic events, the log_10_(number of segments) for the resulting chromosome is drawn from a normal distribution:

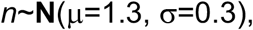

while the log_10_(length) of segments are drawn from a normal distribution

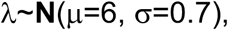

and the start of the chromothriptic event is drawn from a uniform distribution:

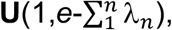

where *e* is the size of the chromosome. The parameters for the distributions were chosen to match the empirical distributions observed in TCGA chromosomes that were called as chromothriptic in the PCAWG dataset.

The breakpoints of the chromothriptic event, [*b_1_*,…,*b_n-1_*], are then the cumulative sums of the segment sizes, apart from the first breakpoint which is 1. The chromosome is then broken into *n* segments by their cumulative lengths, defined by the breakpoints. Whether to lose a segment is drawn from a binomial distribution:

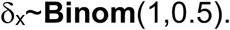

All segments were removed where δ_x_=1. The remaining segments were then randomly reversed if:

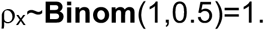

Lastly, the remaining segments were resampled without replacement so that their order is randomized, and are then concatenated together. The chromothriptic chromosome replaces the original chromosome that it originates from.

#### Genome doubling and chromosomal gains/losses

All chromosomes in the set of chromosomes are duplicated to simulate genome doubling. For chromosomal gains, a single chromosome is duplicated, whereas for chromosomal losses a single chromosome is removed.

#### Calculating copy number

Once an assortment of chromosomes has been simulated from a mixture of the previously described processes, the combined copy number across all derivative chromosomes must be calculated across the reference genome. For each reference chromosome, *x*, all segments across the derivative chromosomes that derive from *x* are collated, and the breakpoints across *x* are defined as the ordered unique set of start or end positions of those segments. Then the copy number for segment *i_x_*, is calculated for each allele separately; the A allele copy number is the count of A allele segments in all derivative chromosomes that overlap the segment defined between *b_i,x_* and *b_i+1,x_*, and similar for the B allele copy number. Combined across all reference chromosomes, this gives an allele-specific copy number profile.

#### Combinations of simulations

The following simulations were performed, for 100 samples each:

- CIN×10 – 10 random gain or loss events.
- CIN×50 – 50 random gain or loss events.
- CIN×10->WGD – 10 random gain or loss events, followed by WGD.
- CIN×50->WGD – 50 random gain or loss events, followed by WGD.
- CIN×5->WGD->CIN×50 - 5 random gain or loss events, followed by WGD, followed by 50 random gain or loss events.
- CIN×5->WGD->CIN×25->WGD->CIN×25 – 5 random gain or loss events, followed by WGD, followed by 25 random gain or loss events, followed by WGD, followed by 25 random gain or loss events.
- Chromo. – Chromothripsis of a random chromosome.
- Chromo.->WGD – Chromothripsis of a random chromosome, followed by WGD.
- Chromo.->Amp. – Chromothripsis of a random chromosome, followed by chromosomal gain of the derivative chromothriptic chromosome.
- Chromo.->Amp.->WGD - Chromothripsis of a random chromosome, followed by chromosomal gain of the derivative chromothriptic chromosome, followed by WGD.
- Chromo.->Amp.×5->WGD. Chromothripsis of a random chromosome, followed by chromosomal gain of the derivative chromothriptic chromosome five times, followed by WGD.

For random gain/loss events, a binomial draw was used to decide whether a gain or loss occurred, with p_gain_=0.4.

