## Supplementary Figures for "Signatures of copy number alterations in human cancer"

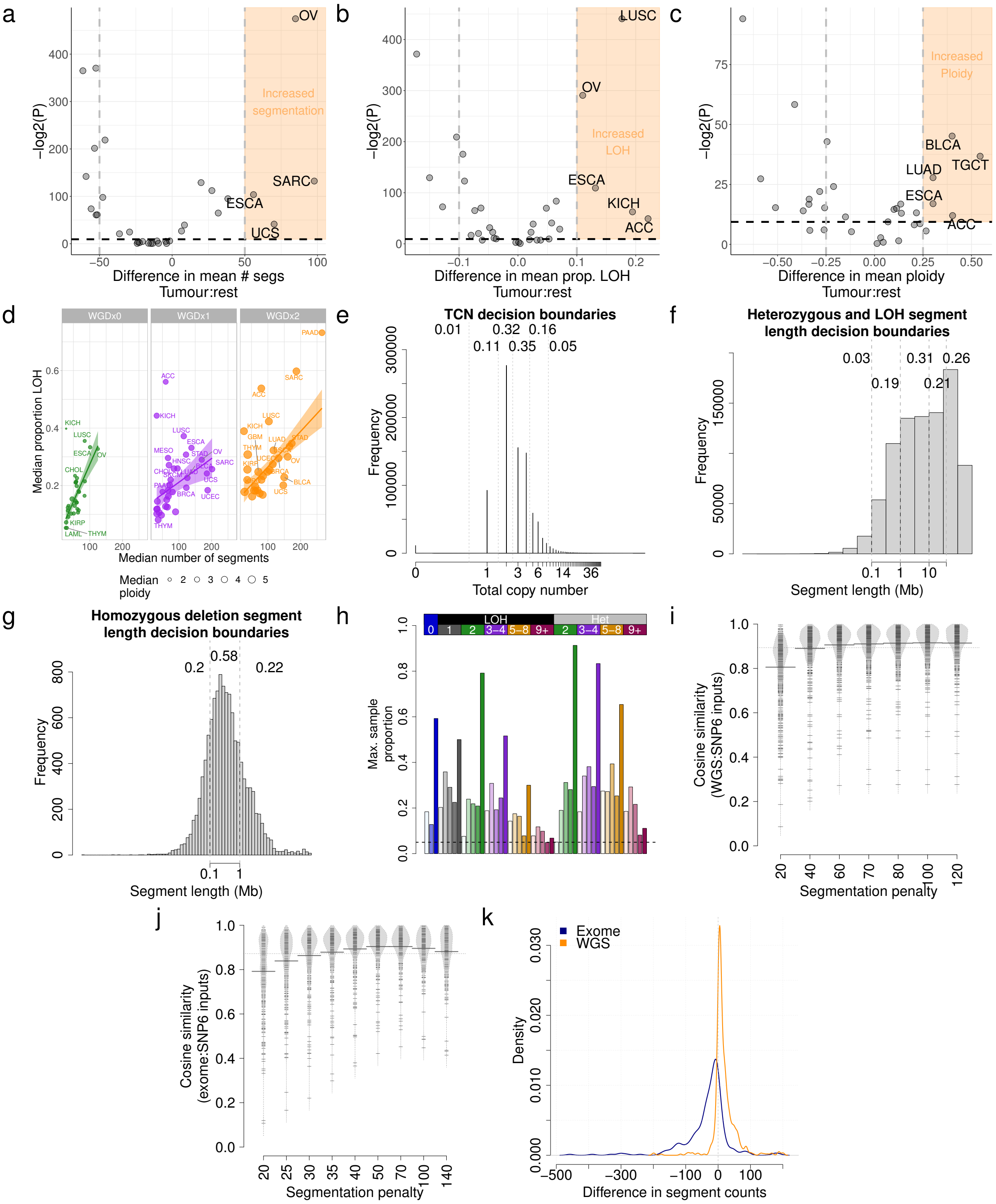

Supplementary Figure 1. Choice of copy number categories.

a) Enrichment of segment counts in TCGA tumour types: x-axis=difference in mean segment counts between tumour type and all other tumours, y-axis=-log2(p-value) from a Mann-Whitney test.
b) Enrichment of LOH in TCGA tumour types: x-axis=difference in mean proportion of genome LOH between tumour type and all other tumours, y-axis=-log2(p-value) from a Mann-Whitney test.
c) Enrichment of high ploidy in TCGA tumour types: x-axis=difference in mean ploidy between tumour type and all other tumours, y-axis=-log2(p-value) from a Mann-Whitney test.

d) Relationship between median number of segments (x-axis), median proportion of the genome that is LOH (y-axis) and ploidy (size) of 33 cancer types in TCGA, split by genome doubling status (panels).
e) Distribution of total copy number across TCGA. Dashed lines indicate decision boundaries between copy number classes. Numbers indicate the proportion of segments across TCGA that fall within the designated category.
f) Distribution of segment lengths across TCGA. Dashed lines indicate decision boundaries between copy number classes. Numbers indicate the proportion of segments across TCGA that fall within the designated category.
g) Distribution of segment lengths for all homozygous deletion (CN=0) segments across TCGA. Dashed lines indicate decision boundaries between copy number classes. Numbers indicate the proportion of segments across TCGA that fall within the designated category.
h) Maximum proportion of segments (y-axis) of each copy number category (x-axis) in any sample across TCGA. Increasing colour saturation indicates increasing segment length.
i) Cosine similarity (y-axis) between input copy number summary vectors for whole genome sequencing and SNP6 array derived copy number profiles.
j) Cosine similarity (y-axis) between input copy number summary vectors for exome sequencing and SNP6 array derived copy number profiles.
k) Difference in segment counts between SNP6 array copy number profiles and whole genome sequencing (orange) or exome sequencing (blue) copy number profiles.

1)

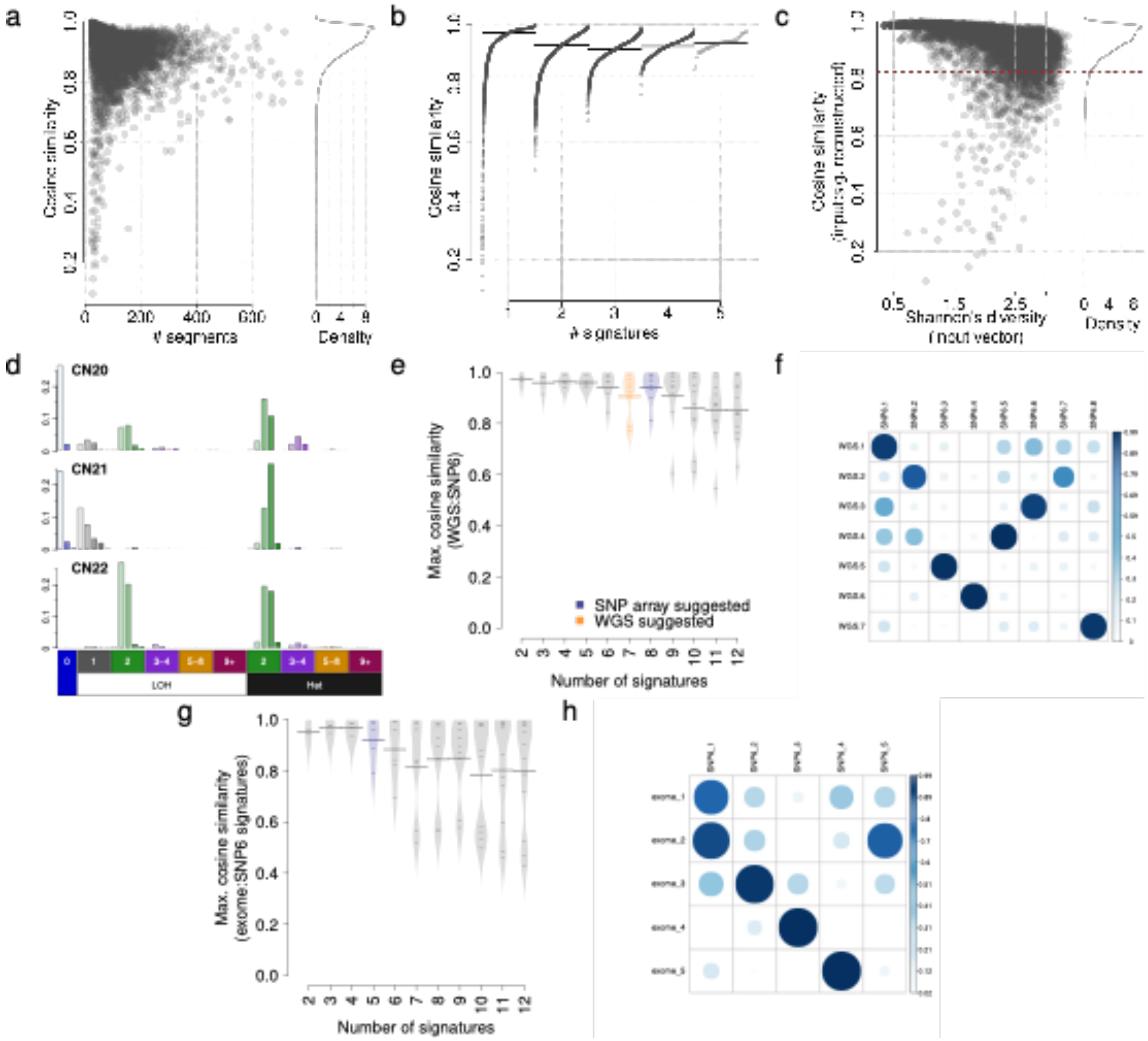


Supplementary Figure 2. Signature derivations

a) Cosine similarity between input copy number 48 dimensional vectors, and signature reconstructed 48 dimensional vectors (y-axis) against number of segements in each copy number profile (x-axis). Dashed line indicates cosine similarity threshold for non-random similarity (p<0.05).
b) Cosine similarity between input copy number 48 dimensional vectors, and signature reconstructed 48 dimensional vectors (y-axis) against the number of signatures assigned in each sample (x-axis). Dashed line indicates cosine similarity threshold for non-random similarity (p<0.05). Solid lines indicate median cosine similarity. The number of signatures is plotted offset by the quantile of the sample.
c) Cosine similarity between input copy number 48 dimensional vectors, and signature reconstructed 48 dimensional vectors (y-axis) against the Shannon's diversity of copy number states in input 48 dimensional vector (y-axis). Dashed line indicates cosine similarity threshold for non-random similarity (p<0.05).
d) Three artefactual signatures identified in the TCGA pan-cancer analysis. Artefactual signatures are typified by a large number of homozygous deletions (top two), or small segment sizes of equal copy number in LOH and heterozygous segments (bottom).

e) Maximum cosine similarities between each WGS signature and any SNP6 identified signatures (i.e. closest matching signature cosine similarity, y-axis) from 512 samples, with varying numbers of signatures decomposed (x-axis).
f) Cosine similarities between WGS (x-axis) and SNP6 (y-axis) identified signatures from 512 samples, with a segmentation penalty of 70.
g) Maximum cosine similarities between each exome signature and any SNP6 identified signatures (i.e. closest matching signature cosine similarity, y-axis) from 282 samples, with varying numbers of signatures decomposed (x-axis).
h) Cosine similarities between exome and SNP6 identified signatures from 282 samples, with a segmentation penalty of 70 and suggested number of signatures extracted.

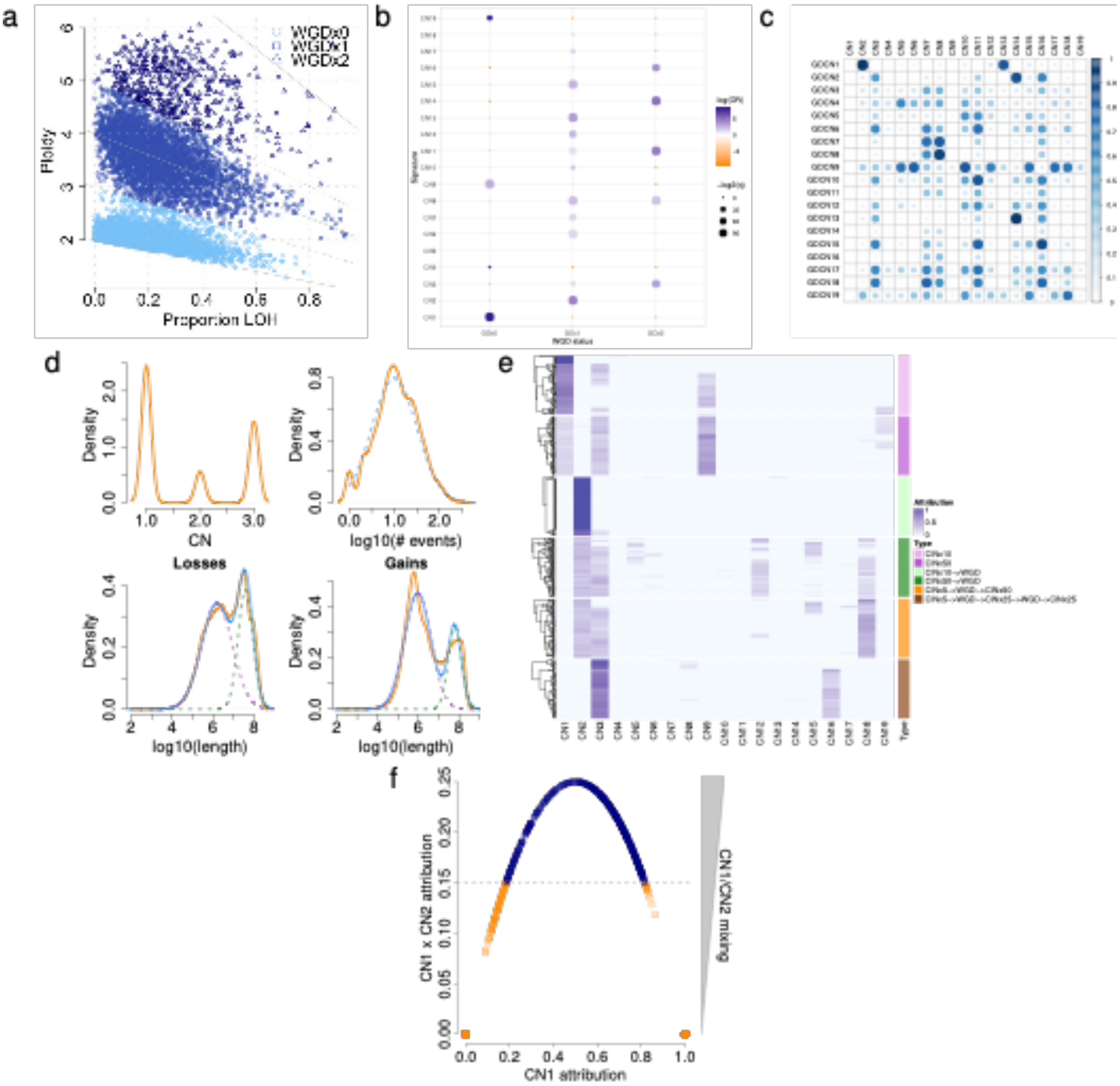


Supplementary Figure 3. Ploidy associated signatures.

a) WGD calls for TCGA, based on ploidy and the proportion of the genome that is LOH.
b) Associations between copy number signature exposure and WGD calls. GDx0=non-genome doubled, GDx1=genome doubled once, GDx2=twice genome doubled.
c) Cosine similarities between signatures (CN1-19) and their artificially genome doubled counterparts (GDCN1-19). A high cosine similarity between e.g. CN2 and GDCN1 indicated that CN2 is a genome doubled version of CN1.
d) Distributions of total copy number of segments (only TCN1-3, top-left), number of non-diploid segements (top-right), segment length of losses (TCN=1, bottom-left) and segment length of gains (TCN=3, bottom-right) for predominantly diploid (CN1+9>0.8) profiles in TCGA. Orange lines indicate empirical distributions, non-orange lines indicate simulated distributions. Dashed lines indicate components of mixture distributions, or the distribution for non-mixed distributions. Solid blue lines indicate joint distributions.
e) Attributions (blue) of the 19 pan-cancer signatures (x-axis) in 6 simulation designs each of 100 samples (y-axis). CIN=random sub-chromosomal copy number gain or loss. WGD=whole genome doubling.

f) CN1 attribution (x-axis) against CN1 attribution x CN2 attribution in samples for which CN1+CN2 attribution = 1. Decision boundary for determining highly aneuploid samples is shown in grey. Orange points are taken for further analysis of aneuploidy.

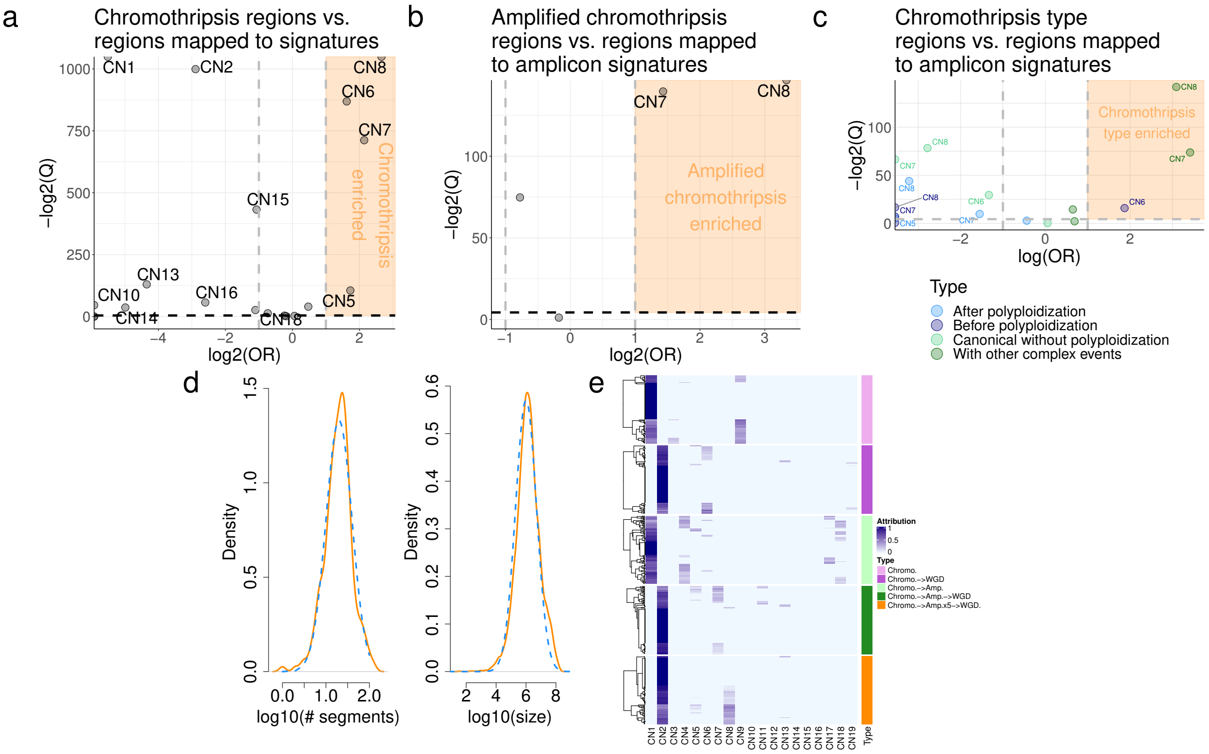


Supplementary Figure 4. Amplicon-associated signatures.

a) Correlation between copy number signature attributed segments and chromothriptic regions at a genomic level. X-axis=effect size (log odds ratio), y-axis=significance (-log2 q-value).
b) Same as for (a), but correlated against amplified chromothripsis. CN7-8 OR= OR=2.69 and 10.05 respectively, q<0.05.
c) Same as for (a), but correlated against distinct chromothripsis types.
d) Distributions of the number of segments (left) and segment sizes (right) on chromothriptic chromosomes identified by PCAWG. Orange lines indicate empirical distributions. Blue dashed lines indicate simulation distributions.
e) Attributions (blue) of the 19 pan-cancer signatures (x-axis) in 5 simulation designs each of 100 samples (y-axis). Chromo.=chromothripsis. WGD=whole genome doubling. Amp=single gain of the derivative chromothriptic chromosome.

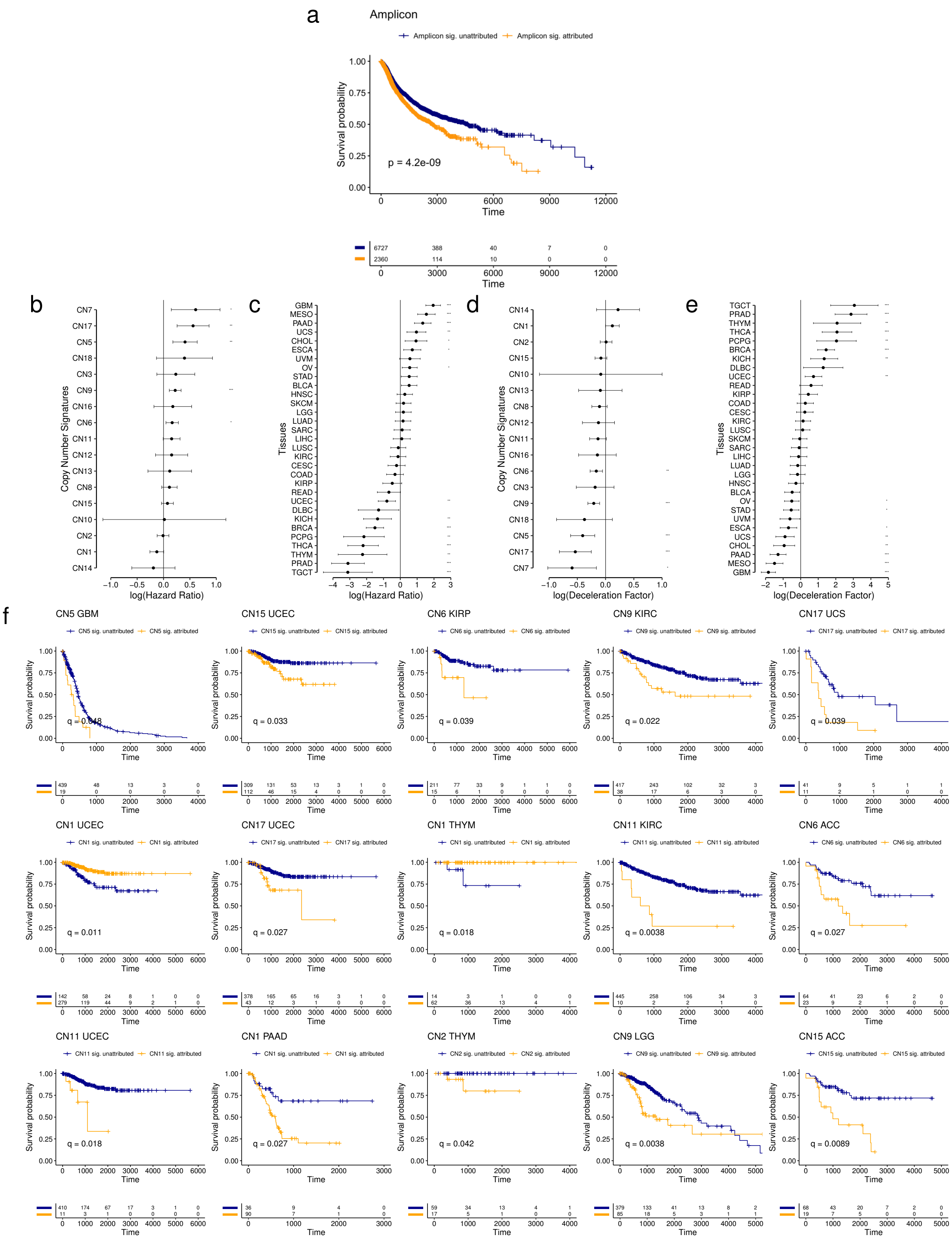

Supplementary Figure 5. Survival associations

a) Kaplan-Meier curves of disease specific survival for patients whose tumours are amplicon signature (CN4:8) attributed (orange) and non-attributed (blue).
b) Cox-model hazard ratios (x-axis) for copy number signatures (y-axis) with copy number signature attribution and tumour type as a covariates (see Supplementary Figure 5e).
c) Cox-model hazard ratios (x-axis) for tumour types (y-axis) with copy number signature attribution (see Supplementary Figure 5d) and tumour type as covariates.

d) Accelerate failure time deceleration factors (x-axis) for copy number signatures (y-axis) with copy number signature attribution and tumour type as a covariates (see Supplementary Figure 5c). A log(deceleration factor)<1 indicates reduced survival time (accelerated failure time), while a log(deceleration factor)>1 indicates increased survival time (deaccelerated failure time).
e) Accelerate failure time deceleration factors (x-axis) for tumour types (y-axis) with copy number signature attribution (see Supplementary Figure 5b) and tumour type as covariates. A log(deceleration factor)<1 indicates reduced survival time (accelerated failure time), while a log(deceleration factor)>1 indicates increased survival time (deaccelerated failure time).
f) Kaplan-Meier curves for within-tumour type associations with copy number signature attribution. Tumour type/copy number signature combinations with a significant effect on survival (q<0.05) are displayed.

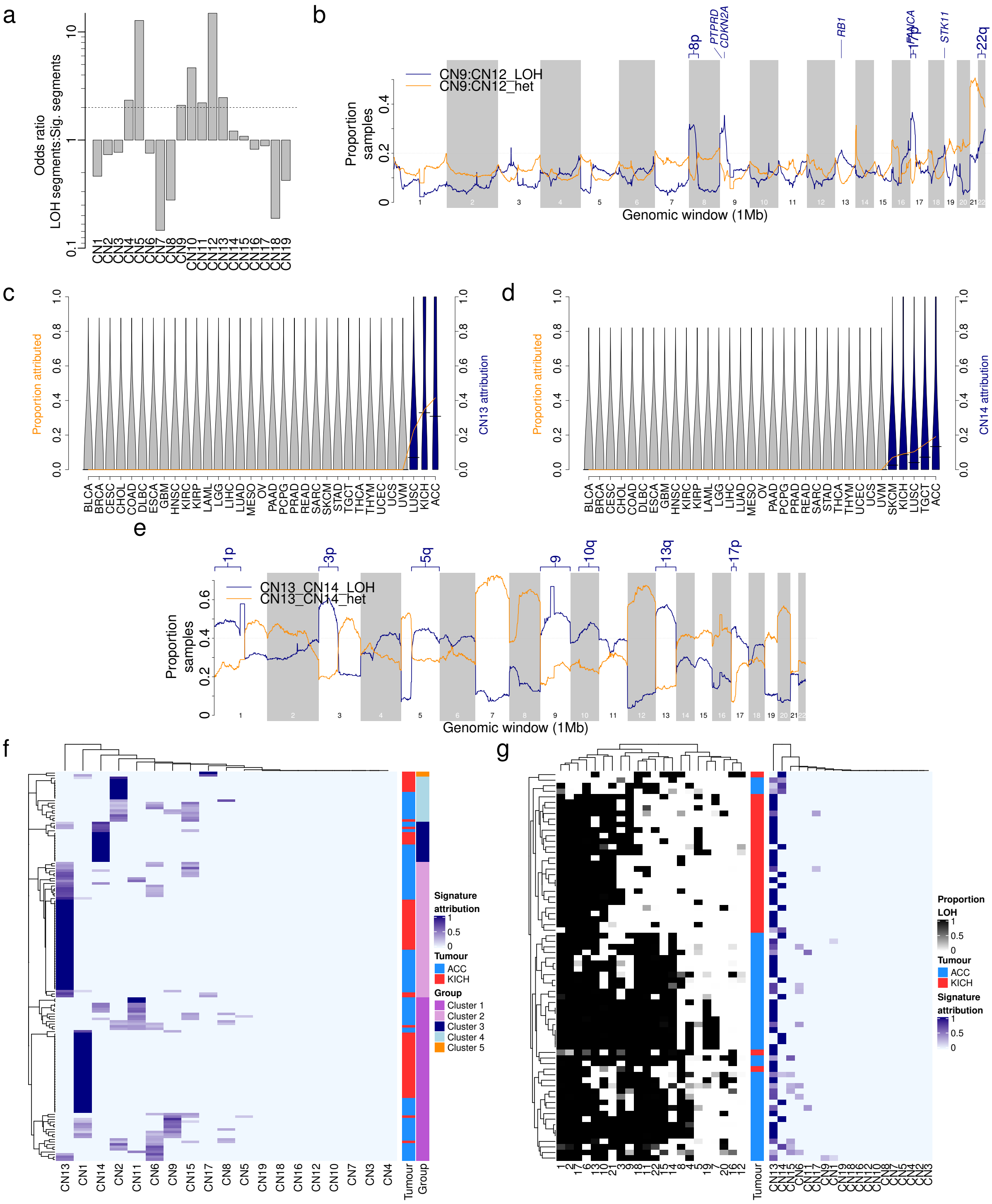


Supplementary Figure 6. LOH associated signatures.

a) Association between LOH segments and mapped copy number signature segments across the full TCGA cohort.
b) Recurrence of mapped LOH signatures (y-axis) across the genome in 1Mb bins (x-axis), split by LOH (blue) or heterozygous (orange) segments. Tumour suppressor genes with >20% of samples with LOH signatures are labelled.
c) Prevalence (orange line) and distribution (violins) of CN13 attributions across TCGA cancer types. Blue violins are cancer types significantly enriched in CN13 compared to all others (q<0.05, Mann Whitney test). KICH enrichment: OR=35, p=1.2e-24, Fisher's exact test. ACC enrichment: OR=51, p=7.4e-43, Fisher's exact test.

d) Prevalence (orange line) and distribution (violins) of CN14 attributions across TCGA cancer types. Blue violins are cancer types significantly enriched in CN14 compared to all others ($q<0.05$, Mann Whitney test). KICH enrichment: OR=8, p=1.9e-4, Fisher's exact test.
e) Recurrence of mapped arm-level LOH signatures (y-axis) across the genome in 1Mb bins (x-axis), split by LOH (blue) or heterozygous (orange) segments. Chromsome arms with >50% of samples with LOH signatures are labelled.
f) Heatmap of signature (x-axis) attributions in ACC and KICH samples (y-axis). Samples are clustered according to their signature attributions (right annotation).
g) Left: Heatmap of LOH prevalence by chromosome (x-axis) and sample (y-axis) for all CN13/CN14 enriched samples (see Supplementary Figure 6f, clusters 2 and 3). Samples are clustered according to chromosomal LOH levels. Right: Copy number signature attributions for the same samples.

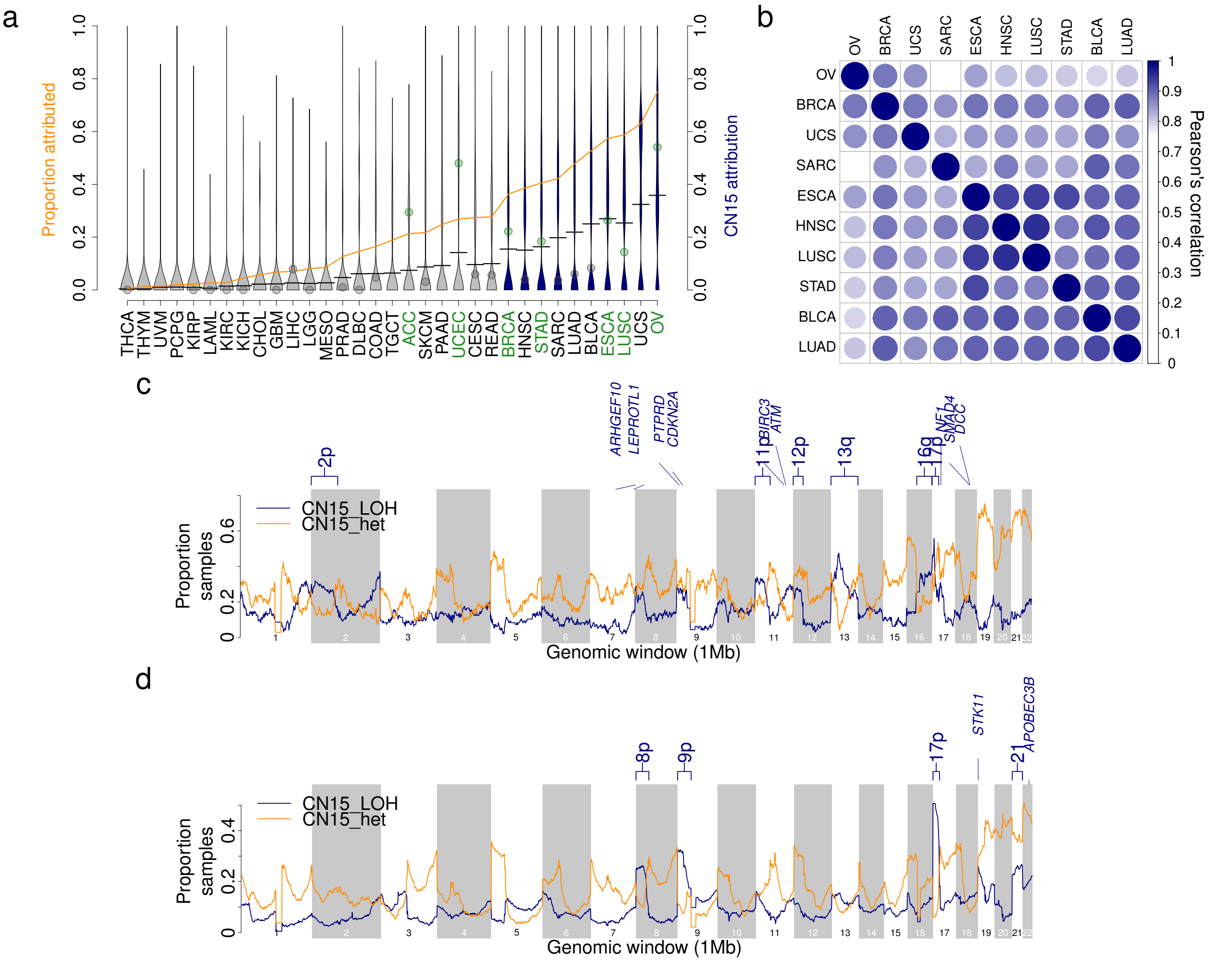


Supplementary Figure 7. Signature of tandem duplicator phenotype

a) Prevalence (orange line) and distribution (violins) of CN15 attributions across TCGA cancer types. Blue violins are cancer types significantly enriched in CN15 compared to all others (q<0.05, Mann Whitney test). Points indicate the prevalence of TDP in given tumour types from the literature (Menghi et al., 2018) coloured by over- (green) or underrepresentation (gray).

b) Pearson's correlation of recurrence of mapping of CN15 to the genome from pairwise comparisons of CN15 enriched tumour types for heterozygous segments.

c) Recurrence of mapped CN15 in 1Mb windows of the human genome in all CN15 attributed SARC samples, split by LOH (blue) and heterozygous segments (orange). Tumour-suppressor genes in regions with >20% samples attributed to CN15 with LOH segments are labelled.

d) Recurrence of mapped CN15 in 1Mb windows of the human genome in all CN15 attributed STAD, LUAD, BLCA, HNSC, ESCA and LUSC samples, split by LOH (blue) and heterozygous segments (orange). Tumour-suppressor genes in regions with >20% samples attributed to CN15 with LOH segments are labelled.

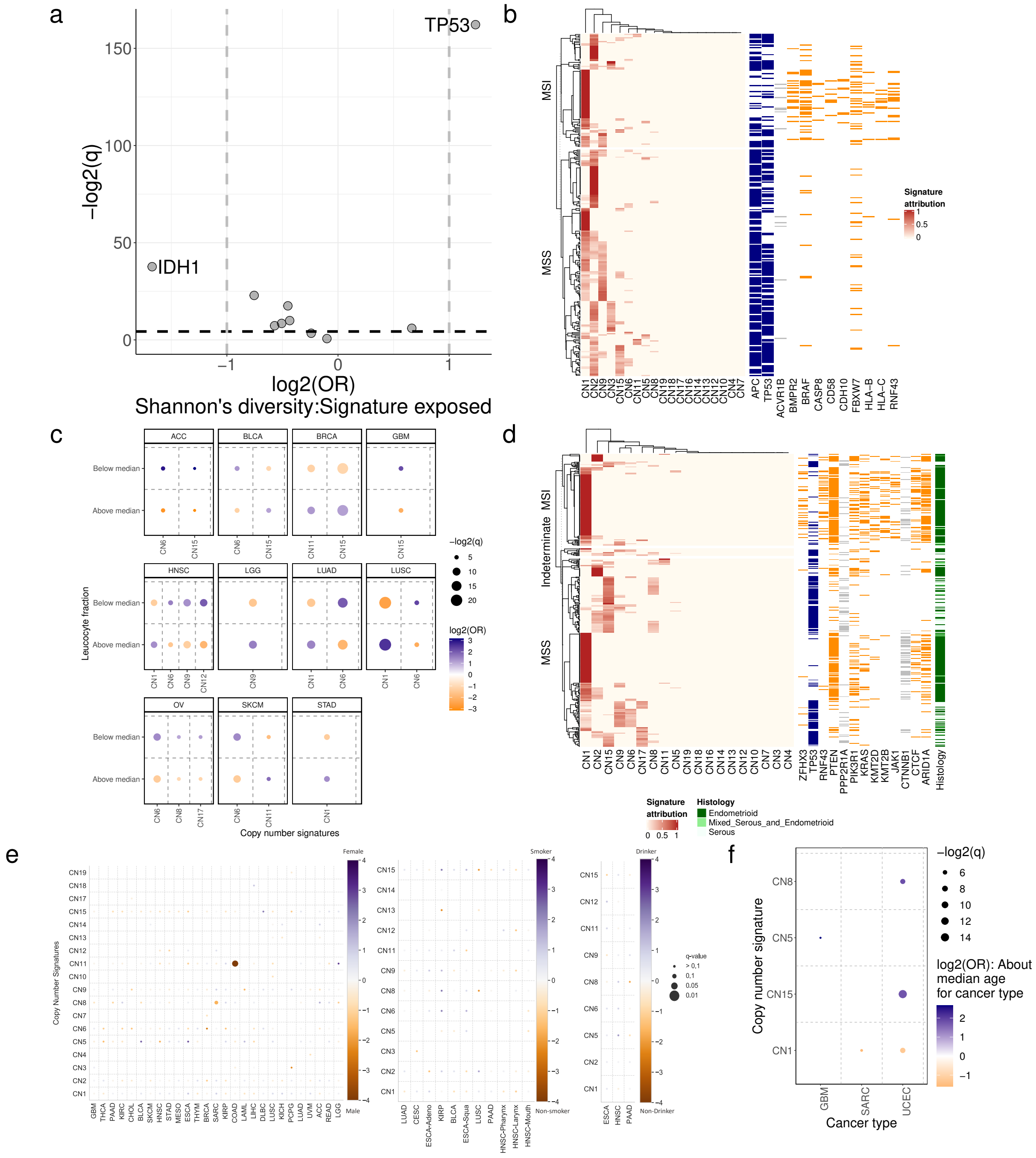


Supplementary Figure 8. Genomic and clinical correlates

a) Correlation between Shannon's diversity index of signature proportions in samples, and driver gene mutation status. Effect size (log2 odds ratio, y-axis) and significance (-log2 q-value, y-axis) are displayed. Driver genes with |log2(OR)|>1 and q<0.05 are labelled. TP53 association: OR=3.42, q=1.5e-49.
b) Heatmap of copy number signature attribution (left) and driver gene mutation status (right) for all COAD samples, split by microsatellite instability status. Driver gene mutations are coloured orange or blue for genes that are positively (OR>1, q<0.05) or negatively (OR<1, q<0.05) associated with MSI status respectively, and grey for genes that are not associated with MSI status (q≥0.05). Association between CN1 or CN2 and MSI status: OR=0.57 and 4.36, p=0.040 and 1.8e-8 respectively, Fisher's exact test.

c) Correlations between leucocyte fraction (y-axis, split by median value per tumour type) and copy number signature attribution (x-axis). Effect size given as log2(OR) (colour) and significance given as q-values (size) are displayed. Only associations with |log2(OR)|>1 and q<0.05 are shown.

d) Heatmap of copy number signature attribution (left) and driver gene mutation status (right) for all UCEC samples, split by microsatellite instability status. Driver gene mutations are coloured orange or blue for genes that are positively (OR>1, q<0.05) or negatively (OR<1, q<0.05) associated with MSI status respectively, and grey for genes that are not associated with MSI status (q≥0.05). Association between CN1 or CN2 and MSI status: OR=0.18 and 2.28, p=2.4e-10 and 2.5e-3 respectively, Fisher's exact test.

e) Correlation between copy number signature (x-axis) attribution and sex (left), smoking status (middle) and drinking status (right) across TCGA samples. Strength of correlation is indicated by colour (orange=anti-correlated, blue=correlated), q-value is indicated by size of point.

f) Association between copy number signatures (y-axis) and median dichotomised age at diagnosis for individual cancer types (x-axis). Strength of correlation is indicated by colour (orange=negatively associated, blue=positively associated), q-value is indicated by size of point. Only tumour types/copy number signature combinations with a significant (q<0.05) association with age at diagnosis are shown.
